# ‘Resistance is futile’: Weaker selection for resistance during larger epidemics further increases prevalence and depresses host density

**DOI:** 10.1101/2021.05.25.445183

**Authors:** Jason C. Walsman, Meghan A. Duffy, Carla E. Cáceres, Spencer R. Hall

**Affiliations:** University of Pittsburgh; University of Michigan; University of Illinois Urbana-Champaign; Indiana University

## Abstract

What determines how much resistance hosts evolve? One might intuit that hosts evolve higher resistance when parasites are more abundant. However, the opposite pattern can arise due to costs of resistance. Here we illustrate with mathematical, experimental, and field approaches how ecological context can increase parasite abundance and select for lower resistance. ‘Resistance is futile’ when all host genotypes become sufficiently infected. To make this argument, we first analyzed an eco-evolutionary model of parasites, hosts, and hosts’ resources. We determined eco-evolutionary outcomes for resistance (mathematically, transmission rate) and densities along gradients that drive epidemic size. When epidemic drivers are high, hosts evolve lower resistance, amplifying epidemics and decreasing host density. Experimental mesocosms qualitatively agreed. In the experiment, higher supply of nutrients drove larger epidemics of survival-reducing fungal parasites. Evolving zooplankton hosts were *less* resistant at high nutrients than at low. Less resistance, in turn, was associated with higher infection prevalence and lower host density. We also analyzed the size of naturally occurring epidemics, finding a broad, bimodal distribution of epidemic sizes consistent with the eco-evolutionary model. Together, our three approaches supported predictions that high epidemic drivers lead to evolution of lower resistance which drives higher prevalence and lower host density.

## Introduction

Epidemics of infectious disease threaten many populations (Dobson et al. 2008), including posing risks to livestock (Horan and Fenichel 2007) and species of conservation concern, such as birds and amphibians (Cooper et al. 2009; Vredenburg et al. 2010). Parasites harm their hosts (“virulence”) by increasing mortality (“mortality virulence”) or decreasing fecundity (“fecundity reduction or castration”: Ebert et al. 2000), reducing host density and increasing extinction risk (Ebert et al. 2000). Because parasites lower fitness, and because hosts vary in their resistance (specifically preventing infection, one resistance pathway), we expect resistance to evolve in host populations during parasite outbreaks. If this evolution of resistance occurs quickly enough, it should reduce the size of epidemics and, consequently, their impact on host density (Penczykowski et al. 2011). Thus, rapid evolution of increased resistance by hosts is often expected to suppress prevalence and prevent declines of host density (Altizer et al. 2003; Christie and Searle 2018; Duffy and Sivars-Becker 2007).

However, the evolutionary response of hosts during epidemics can hinge upon costs of resistance. Resistance becomes costly when linked to fecundity or other traits (Auld et al. 2013; Boots and Begon 1993; Duncan et al. 2011; Hall et al. 2010; van der Most et al. 2011). Given these costs, host populations should evolve decreased resistance to infection during times without parasites. Such evolution occurs for various hosts, e.g., poultry (van der Most et al. 2011), wildflowers (O’Hara et al. 2016), paramecia (Duncan et al. 2011) and nematodes (via evolution of selfing: Slowinski et al. 2016). Additionally, hosts can evolve less resistance when epidemics remain small (Duffy et al. 2012). However, most work expects that higher parasite density should grant fitness advantages to costly resistance (Boots et al. 2009; Duffy and Forde 2009; Duffy et al. 2012; Frank 1994; Koskella 2018; Lopez-Pascua et al. 2014). This prediction holds especially strongly for parasites that eliminate reproduction of infected hosts, a severe form of virulence (Bohannan and Lenski 1997; Bohannan and Lenski 2000; Boots and Haraguchi 1999; Gomez et al. 2015; Lopez-Pascua et al. 2014). Thus, despite costs of resistance, models typically predict that larger epidemics of virulent parasites should more strongly select for resistance. Such trait evolution should lower prevalence, thereby reducing depression of host density during epidemics.

However, there is another possibility. During large epidemics, low resistance with high reproduction might provide the fittest host strategy. This response can arise when infected hosts can still reproduce (Best et al. 2017; Bonds et al. 2005; Donnelly et al. 2015; Miller et al. 2007). With some fecundity retained, low resistance-high fecundity genotypes become most fit when parasites infect all genotypes. More generally, the most fit resistance-fecundity strategy becomes a hump-shaped function of epidemic size (Donnelly et al. 2015). Scarce parasites cause evolution of low resistance because the fitness benefits of higher fecundity outweigh survival benefits of resistance. At intermediate density of parasites, the survival benefits of higher resistance begin to outweigh benefits of higher fecundity (Donnelly et al. 2015). Then enters the new(er) insight: at even higher parasite density, lower resistance becomes the fitter strategy again. Because prevalence is high enough, the survival benefits of resistance contribute less (Donnelly et al. 2015). Instead – as long as infected hosts can still reproduce – the benefits of higher fecundity reign supreme. In this scenario, during large epidemics, evolution of higher resistance does little to avoid virulent effects on mortality and/or fecundity. Instead, hosts with low resistance reap the benefits of increased fecundity (Donnelly et al. 2015). Hence, when risk infection becomes high enough, ‘resistance is futile’ (a shorthand for a peripherally similar purpose also used by Best et al. 2010).

We illustrate conditions and consequences for ‘resistance is futile’ evolution during epidemics with math and data. Like in previous work (e.g., Donnelly et al. 2015), we calculate the adaptive dynamics of host evolution using a mathematical proxy for host fitness (an eigenvalue from a next generation matrix). To aid transparent interpretation, we also decompose selection into fecundity and mortality components of fitness. This direct fitness approach demonstrates that resistance only slightly decreases prevalence when prevalence is high, the mechanism for ‘resistance is futile.’ The same logic applies when ecological factors (‘epidemic drivers’, e.g., resources for hosts) push epidemics to be large. As an important next step, we demonstrate the population-level outcomes from host evolution for infection prevalence and host density, unlike previous work (see Miller et al. 2007 for a partial exception). To contextualize those eco-evolutionary outcomes, we compare them to non-evolving, ecology-only populations (like used in our experiment), and we create ‘ecological derivative maps’. This contextualization shows that high epidemic drivers can lead to evolution of lower resistance. That response of hosts increases infection prevalence and lowers host density.

We test the model’s predictions qualitatively with a mesocosm experiment and a field survey with a planktonic system. In this system, infection by the fungal parasite (*Metschnikowia bicuspidata*) increases mortality but infected zooplankton hosts (*Daphnia dentifera*) can still reproduce. Additionally, hosts face a fecundity-resistance tradeoff (via a feeding rate mechanism: Auld et al. 2013; Hall et al. 2010). We manipulated an epidemic driver: higher supply of nutrients for host’s resources supported denser host populations and larger epidemics. All populations started with the same mean level of resistance. After clonal turnover, high nutrient (high parasite) populations were less resistant than low nutrient (low parasite) populations. Across populations, lower resistance was associated with higher prevalence and lower host density, consistent with the model. Field support from this prediction arose in a survey of naturally occurring fungal epidemics in lakes. In this survey we found a somewhat bimodal and right-skewed distribution of epidemics with higher variance. This pattern echoed that produced by positive feedbacks in the eco-evolutionary model. Thus, the model, experiment, and natural populations all support a ‘resistance is futile’ effect. These responses raise concern that host evolution during large, virulent outbreaks can lead to larger outbreaks that depress density.

### Eco-evolutionary model of resources, hosts, and parasites

The ‘resistance is futile’ prediction arises from an eco-evolutionary model analyzed with Adaptive Dynamics. Inspired by the planktonic system, this model allows for full feedbacks of resources (*R*), susceptible hosts of genotype i (*S*_i_), infected hosts (*I*_i_), and free-living parasite propagules (*Z*). We use reasonable parameters values for the planktonic system (Table 1):

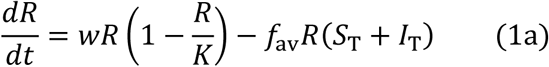

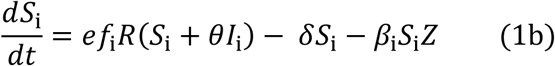

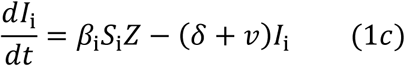

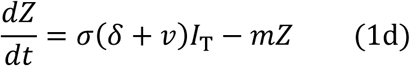

**Table 1.** Symbols, meaning, and default values or ranges for state variables (top), functions of state variables (middle), and traits and other parameters (bottom) in an eco-evolutionary model of hosts, disease, and resources (eq. 1).

| Symbol | Meaning | Units | Value or range |
| --- | --- | --- | --- |
| $t$ | Time | day | Varies |
| $R$ | Density of resources | $\mu\text{g chl } a \cdot \text{L}^{-1}$ | See figures |
| $S_i$ | Density of susceptible hosts $i$ | $\text{hosts} \cdot \text{L}^{-1}$ | See figures |
| $I_i$ | Density of infected hosts $i$ | $\text{hosts} \cdot \text{L}^{-1}$ | See figures |
| $Z$ | Density of parasite propagules | $\text{parasites} \cdot \text{L}^{-1}$ | Varies |
| $H_i$ | Total density of hosts $i$, $S_i + I_i$ | $\text{hosts} \cdot \text{L}^{-1}$ | See figures |
| $p_i$ | Prevalence of infection, $\frac{I_i}{S_i+I_i}$ | unitless | See figures |
| $dp_i/d\beta_i$ | Differential prevalence | parasites·day·L<sup>-1</sup> | See figures |
| $b_i$ | Per-capita fecundity, hosts: $b_i = ef_i(\beta_i)R(1-p_i+\theta p_i)$, eq. 3b | day<sup>-1</sup> | Varies |
| $db_i/d\beta_i$ | Differential fecundity | parasites·L<sup>-1</sup> | See figures |
| $d_i$ | Per-capita mortality, hosts: $d_i = \delta + p_i v$, eq. 3b | day<sup>-1</sup> | Varies |
| $dd_i/d\beta_i$ | Differential mortality | parasites·L<sup>-1</sup> | See figures |
| $r_i$ | Fitness, hosts: $r_i = b_i - d_i$ | day<sup>-1</sup> | Varies |
| $dr_i/d\beta_i$ | Differential fitness | parasites·L<sup>-1</sup> | See figures |
| $f_{\max}$ | Maximum feeding rate | L·host<sup>-1</sup>·day<sup>-1</sup> | 0.0039 |
| $f_{\min}$ | Minimum feeding rate | L·host<sup>-1</sup>·day<sup>-1</sup> | 0.0024 |
| $f_{\text{scale}}$ | Tradeoff parameter, eq. 2 | parasite·host<sup>-1</sup> | 1 x 10<sup>4</sup> |
| $e$ | Conversion efficiency | hosts· μg chl *a*<sup>-1</sup> | 0.25* |
| $f_i$ | Feeding rate, host i | L·host<sup>-1</sup>·day<sup>-1</sup> | 3.24-3.9 x 10<sup>-3†</sup> |
| $K$ | Carrying capacity, resource | μg chl *a* ·L<sup>-1</sup> | 100, 52-120* |
| $m$ | Loss rate of parasites | day<sup>-1</sup> | 0.5* |
| $v$ | Mortality virulence | day<sup>-1</sup> | 0.037* |
| $w$ | Intrinsic rate of increase, $R$ | day<sup>-1</sup> | 0.9* |
| $\delta$ | Background mortality rate | day<sup>-1</sup> | 0.05* |
| $\beta_i$ | Transmission rate from propagule to host i | L·parasite<sup>-1</sup>·day<sup>-1</sup> | 0.334-9.75 x 10<sup>-6*</sup> |
| $\theta$ | Fecundity of infected relative to susceptible hosts. | Unitless | 0 (castration) – 1 (full fecundity) |
| $\sigma$ | Parasites released per host | parasites·host<sup>-1</sup> | 100,000* |
| *Biologically reasonable values of traits (Strauss *et al.* 2015) |  |  |  |
| †Biologically reasonable during an epidemic (Strauss et al. 2019) |  |  |  |

Resources grow logistically with intrinsic rate of increase *w* and carrying capacity *K* (first term, eq. 1a). They are consumed by the total sum of susceptible (*S*_T_) and infected (*I*_T_) hosts of genotypes i = 1 to n 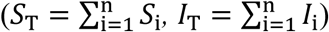 feeding at weighted average rate, *f_av_* 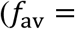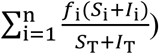 (second term, eq. 1a). Each host genotype i has fixed trait values, the transmission rate of infection from the environment to hosts *β*_i_ and feeding rate *f*_i_. Here, resistance takes the form of decreased transmission rate, an “avoidance” form of resistance that prevents infection (Boots and Bowers 1999); resistance can function by avoiding exposure and/or reducing the chance of infection given exposure. Through feeding, hosts also compete for their shared resources (*R*: eq. 1a). Both host classes, *S*_i_ and *I*_i_, may convert these resources into susceptible offspring (i.e., transmission is purely horizontal; first term, eq. 1b). Susceptible hosts do so with a conversion efficiency *e*. Meanwhile, infected hosts do so with modified efficiency *eθ*, where *θ* = 1 means full fecundity while *θ* = 0 is castration (no offspring production). Outcomes for intermediate values of *θ* are highlighted briefly (see Fig. 1B-D; dashed line) and presented more fully elsewhere (Fig. A1). Genotypes with higher feeding rate *f*_2_ > *f*_1_ have higher fecundity (*ef*_2_*R* > *ef*_1_*R* if uninfected or *ef*_2_*Rθ* > *ef*_1_*Rθ* if infected). All genotypes experience identical background mortality rate *δ* (second term, eq. 1b). Additionally, susceptible hosts can become infected at rate *β*_i_ by encountering parasite propagules while feeding (third term, eq. 1b).

**Figure 1.**
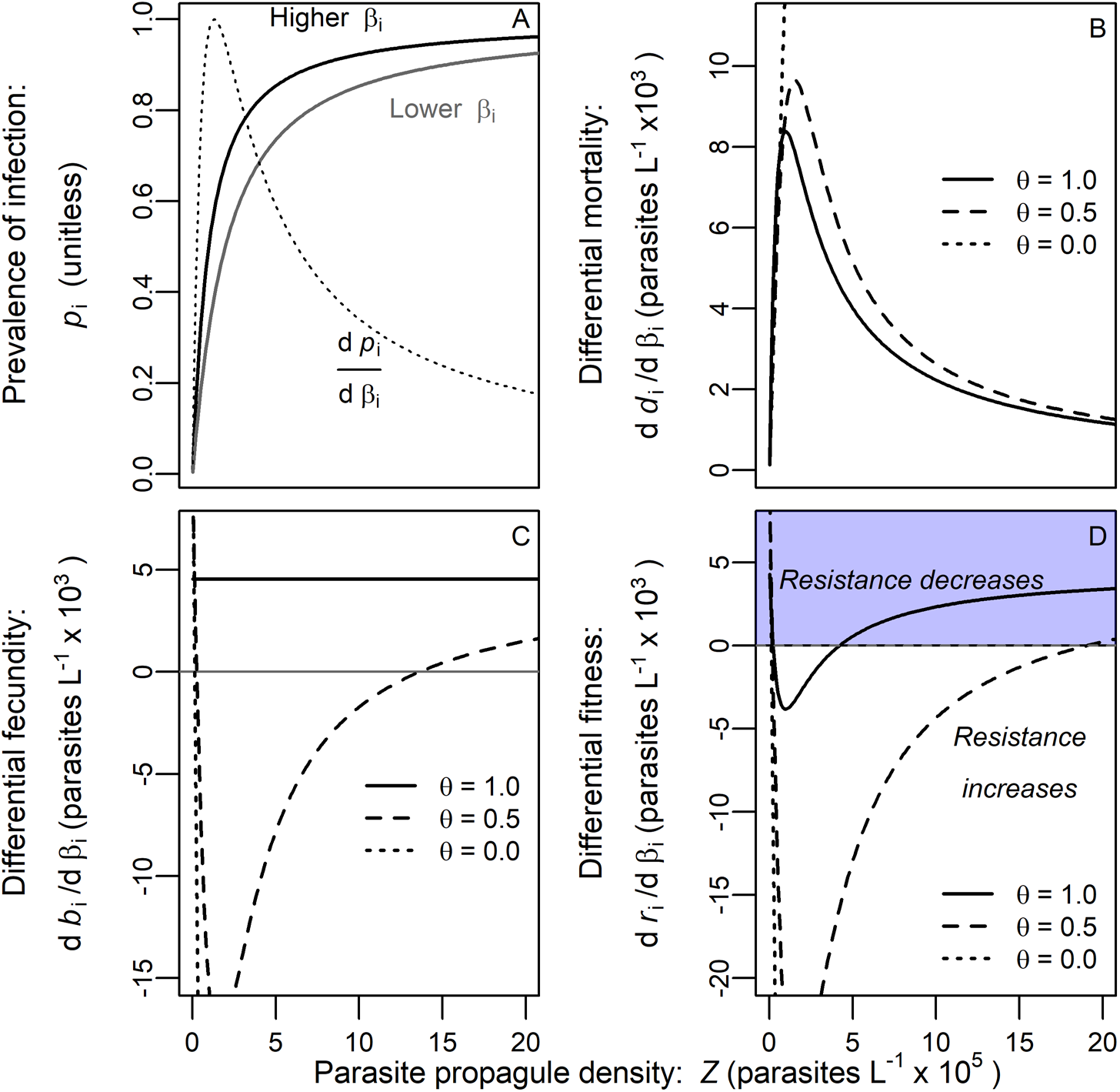
A direct fitness approach to modeling selection on resistance at fixed density of parasite propagules (Z) and resources (R). (A) As parasite propagules increase (higher *Z*, x-axis), prevalence (*p*_i_) saturates, faster for a high transmission rate genotype (black line, *β*_i_ = 1.15 x 10^-6^) than a low one (gray, *β*_i_ = 5.75 x 10^-7^). Differential prevalence (the marginal effect of transmission rate on prevalence, d*p*_i_/d*β*_i_; tied to vertical distance between solid black and gray but scaled for graphing purposes to fall between 0 and 1) peaks at intermediate *Z*. (B) Because it is proportional to differential prevalence, differential mortality (d*d*_i_/d*β*_i_: eq. 3b) peaks at intermediate *p*_i_ (but, at *θ* = 0, host extinction occurs first). Dotted line: *θ* = 0, castration; dashed line: *θ* = 0.5, reduced fecundity; solid line: *θ* = 1, full fecundity. (C) Without disease (*Z* = 0), fecundity increases with transmission rate (d*b*_i_/d*β*_i_ > 0, due to the costs of resistance: eq. 2). That benefit remains at *θ* = 1. When *θ* < 1, differential fecundity first decreases with *Z* but then increases (and can become positive if *θ* > 0; gray line d*b*_i_/d*β*_i_ = 0). (D) At low parasite density, differential fitness (d*r*_i_/d*β*_i_) selects for increased transmission rate (decreased resistance; blue region). At intermediate parasite density, selection favors decreasing transmission rate [increased resistance; all three curves below 0]. High parasite density selects for increased transmission rate again if *θ* > 0 (for *θ* = 0.5 and 1, the curves return to positive). *β*_i_ = 1.15 x 10^-6^. *R* was chosen to help offset fecundity reduction for each curve: *θ* = 0, *R* = 250; *θ* = 0.5, *R* = 200; *θ* = 1, *R* = 97.2. Other parameters follow defaults (Table 1).

Feeding and transmission rate are connected by a tradeoff (eq. 2):

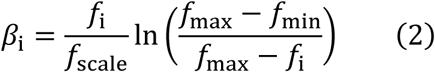

as governed by three positive shape parameters (*f*_max_, *f*_min_, and *f*_scale_) and where ln(…) is the natural logarithm. Hence, transmission rate *β*_i_ can range from zero to infinity, yet feeding rate *f*_i_ remains bounded, ranging from *f*_min_ to *f*_max_ as scaled by a constant (*f*_scale_). With this tradeoff, two points arise. First, it links three host traits: higher feeding rate (*f*_i_) is associated with higher per capita fecundity (*ef_i_R* or *ef_i_Rθ*, at a given *R*) and increased transmission rate *β*_i_ (i.e., decreased resistance). Second, this tradeoff shape imposes accelerating costs: increased resistance (decreased *β*_i_) corresponds to a small cost (decrease *f*_i_) when resistance is low (high *β*_i_). But when resistance is high (low *β*_i_), even further increases of resistance impose very costly feeding reduction. We chose this shape conservatively, to prevent extreme evolution. Accelerating costs typically select for intermediate trait values while decelerating costs tend to select for extreme ones (Boots and Haraguchi 1999). Hence, the asymptotic limit on feeding rate (*f*_max_) prevents the ‘resistance is futile’ effect from selecting for higher and higher feeding rate, without limit. Yet, despite the conservativeness of this tradeoff (eq. 2), hosts can still evolve extremely high or low transmission rates in the model (Figs. 2D, A1A).

**Figure 2.**
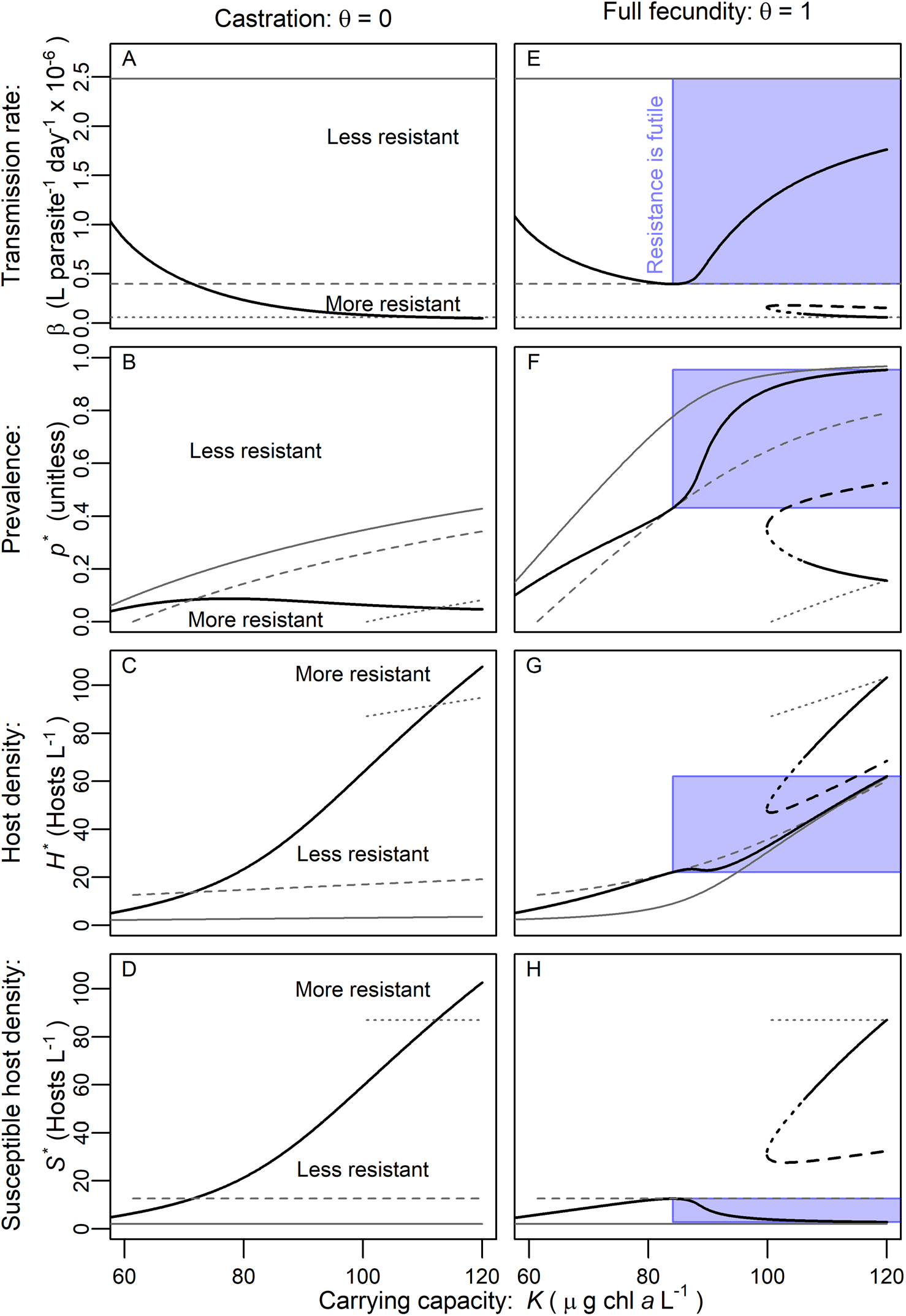
Higher carrying capacity and larger epidemics can cause hosts to evolve less resistance. Solid black curves represent eco-evolutionary outcomes for transmission rate (*β_CSS_*; inverse of resistance) across a carrying capacity gradient (*K*). Gray curves denote response of single reference genotypes (ecology-only populations: high *β*: solid line; medium *β*: dashed; low *β*: dotted). Gray curves only drawn in the range where the endemic equilibrium is feasible. *Castration (θ = 0):* (A) Hosts evolve increasing resistance (decreasing *β*_CSS_) with *K*, which (B) keeps prevalence, *p*\*_CSS_, much lower relative to ecology-only cases [constant *β*; *p*\*_eco_ increases with *K*]. Higher resistance and lower *p*\*_CSS_ allows density of (C) total hosts, *H*\*_CSS_, and (D) susceptible hosts, *S*\*_CSS_ to increase with *K*. *No effect of infection on fecundity (θ = 1):* (E) Higher resistance first evolves with *K*, then lower resistance (in the ‘resistance is futile’ blue box). With higher *K* yet, alternative high or low resistance attractors (black curves) emerge, separated by an unstable evolutionary repeller (dashed black). For a small interval, the lower critical point is an evolutionary branching point (dotted black). (F) Prevalence increases with *K*, more slowly at first, then more quickly as hosts evolve decreasing resistance (blue box). Prevalence quickly becomes very high unless hosts evolve to the more resistant attractor. (G) Total and especially (H) susceptible host density remain low with *K* at the low resistance attractor (but can otherwise increase at the more resistant attractor). See Table 1 for default parameter values.

Infection converts susceptible hosts into infected hosts (first term, eq. 1c). These hosts suffer elevated death rate due to mortality virulence of infection, *δ* + *v* (where *v* is the added mortality assumed equal for all genotypes; second term eq. 1c). When infected hosts die, all genotypes release *σ* parasite propagules, *Z*, back into the environment (first term, eq. 1d). Losses of parasite propagules occur at background rate *m* (second term, eq. 1d).

### Analysis of selection on transmission rate by parasites via a direct fitness approach

#### Methods

To aid interpretation of the core model mechanism, we start with analysis of selection on resistance (*β*) via fitness components of the host. Specifically, we focus on how parasites alter the relationship between *β*_i_ and fitness in a constant environment (without eco-evolutionary feedbacks). Given prevalence of infection (*p*_i_), host fitness (*r*_i_) is the per-capita rate of increase for a given genotype (eq. 3a, b respectively):

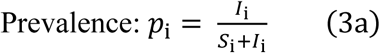

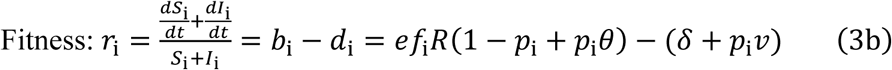

Fitness is the difference of *b*_i_ and *d*_i_ (eq. 3b). The expression for birth rate, *b*_i_ = *ef*_i_*R*(1-*p*_i_*+p*_i_*θ*), denotes the average per-capita birth rate as a function of resources, *R*. The proportion of susceptible hosts (1-*p*_i_) contribute fully to fecundity while the contribution from infected hosts (*p*_i_*θ*) depends on the degree to which infected hosts retain fecundity (expressed by *θ*). The per capita death rate is *d*_i_ = *δ* + *p*_i_*v*. The *p*_i_*v* term increases death rate with prevalence due to mortality virulence. Therefore, fitness depends on resources (*R*) and prevalence of infection (*p*_i_).

Given fitness, *r_i_*, we can calculate selection after evaluating prevalence in a constant environment (set at resources, *R,* and parasite propagules, *Z*). After deriving dynamics of prevalence, d*p*_i_*/*d*t* (using eq. 1b, c), we find a stable value of prevalence, i.e., one with zero change through time (d*p*_i_*/*d*t* = 0) and that is stable [ensured by ∂/∂*p*_i_ (*dp*_i_*/dt*) < 0; see Appendix section 1]. This stable prevalence, *p*_i_(*β*_i_, *R*, *Z*), once expressed in terms of a genotype’s traits (*β*_i_) and its environment (*R* and *Z*), allows calculation of fitness (using eq. 3b) and, importantly here, its birth and death components. This method yields the same fitness optima as produced by the fitness proxy in the more common Next Generation Matrix approach (Diekmann et al. 1990); see Hurford et al. (2009) for clear application of NGM theory to host fitness.

Selection on transmission rate (*β*) then follows from the effect of *β*_i_ on fitness (‘differential fitness’: d*r*_i_/d*β*_i_). It proves illuminating to separate the birth (*b*_i_) from death (*d*_i_) component of fitness (*r*_i_, eq. 4a). Thus, we calculate differential fitness (eq. 4b):

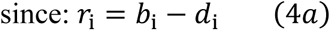

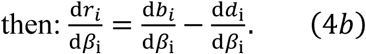

In this expression, differential fitness (d*r*_i_/d*β*_i_) is the difference of differential fecundity (d*b*_i_/d*β*_i_) versus differential mortality (d*d*_i_/d*β*_i_). We calculated these components at different levels of infected fecundity relative to susceptible hosts (*θ*) in a constant *R*-*Z* environment. At these *θ*-*R*-*Z* combinations, if fitness increases with transmission rate (d*r*_i_/d*β*_i_ > 0), then hosts will evolve increased transmission rate (decreased resistance). In contrast, if fitness decreases with transmission rate (d*r*_i_/d*β*_i_ < 0), evolution of lower transmission (higher resistance) ensues.

#### Results

A ‘resistance is futile’ effect arises at high infection prevalence if infected hosts retain some fecundity (*θ* > 0). This result hinges upon the saturation of prevalence (shown for full fecundity, *θ* = 1, but qualitatively similar for other values of *θ*). Prevalence increases with transmission rate (*β*_i_) and density of parasite propagules (*Z*) up to a maximum of 1 (i.e., 100% infection; Fig. 1A). This eventual saturation means that differential prevalence (d*p*_i_/d*β*_i_, marginal increase in prevalence with transmission rate) peaks at intermediate *Z* (here around *p*_i_ = 0.4). Following differential prevalence, differential mortality shows a similarly hump-shaped relationship (because d*d*_i_/d*β*_i_ = *v*\*d*p*_i_/d*β*_i_; Fig. 1B). This pattern does not differ qualitatively with degree of fecundity reduction (all *θ* curves in Fig. 1B). However, castrating (*θ* = 0) parasites would drive host populations extinct before the peak of the hump is reached (i.e., the curve is truncated). Overall, abundant parasites inflict similarly high prevalence on different transmission rates. Therefore, the marginal mortality cost of higher transmission rate shrinks when parasites become highly abundant.

The response of fecundity, *b*_i_, to transmission rate (*β*_i_) depends on the benefits of higher feeding rate (via the tradeoff) vs. benefits of preventing infection that reduces fecundity. With no parasite propagules (*Z* = 0), hosts experience no infection risk. Hence, higher transmission rate only leads to higher fecundity via faster feeding (hence, all d*b*_i_/d*β*_i_ curves start above 0 on the left, Fig. 1C). Then, with full fecundity retained (*θ* = 1), this pattern holds: higher *Z* does not depress differential fecundity (so d*b*_i_/d*β*_i_ stays flat and positive: solid line, Fig. 1C). However, when parasites reduce fecundity (*θ* < 1), higher *β*_i_ genotypes suffer higher prevalence; therefore, a higher proportion of individuals of those genotypes suffer fecundity reduction than individuals of lower *β*_i_ genotypes. Higher density of reproduction-reducing parasites thus penalize high transmission rate genotypes (*θ* = 0.5 curve decreasing, Fig. 1C; see eq. 3b). But with even more parasites, genotypes are similarly infected and suffering fecundity reduction at similar rates (see dashed curve increasing in Fig. 1C). When genotypes are highly infected regardless of *β*_i_, the fecundity benefits of high transmission rate (from tradeoff) exceed those of infection avoidance (dashed curve increases above 0, Fig. 1C). Hence, in environments with low or high enough parasite density, the high transmission-high fecundity strategy is fittest from a fecundity standpoint. But this outcome requires at least some reproduction by infected hosts so that high transmission rate genotypes can be the most fecund when all genotypes are highly infected.

Differential fecundity and mortality combine to govern selection because they determine differential fitness, i.e., the selection gradient. The net effect merely requires subtracting curves (eq. 4b). Flat or U-shaped differential fecundity (d*b*_i_/d*β*_i_; Fig. 1C) minus hump-shaped differential mortality (d*d*_i_/d*β*_i_; Fig. 1B) produces a U-shaped relationship for differential fitness (d*r*_i_/d*β*_i_, Fig. 1D). At low densities of parasite propagules (*Z*), the fecundity advantage of the tradeoff selects for increased transmission rate (*β*; all three curves have d*r*_i_/d*β*_i_ > 0). At intermediate *Z*, mortality (and fecundity reduction if *θ* < 1) selects for decreased *β* (all three curves have negative values). But at high *Z*, differential mortality declines while differential fecundity increases or remains high. Consequently, as long as infected hosts can reproduce (*θ* > 0), the fecundity advantage of high *β* can select for lower resistance at high *Z* (higher *β*; dashed and solid curves in Fig. 1D). Host populations withstand such high parasite densities best if resource density is elevated (as detailed in Appendix: section 1).

This analysis of selection on fitness components provides an intuitive roadmap for the ‘resistance is futile’ outcome during full eco-evolutionary epidemics. When hosts face a resistance-fecundity tradeoff, they have to balance costs and benefits of resistance. If epidemics stay small, low resistance provides fecundity benefits. If epidemics become larger, the survival costs of low resistance become more pronounced. Fecundity reduction (0 < *θ* < 1) only exacerbates the costs of low resistance. Hence, parasites select for resistance. But once parasites become very dense – and epidemics become large – resistance grants diminishing marginal benefits. Instead, as long as infected hosts reproduce, the fecundity benefits of low resistance pre-dominate once again. Assuming hosts populations can withstand such high propagule density, this selection analysis predicts evolution according to a ‘resistance is futile’ response during large epidemics.

### Full eco-evolutionary outcomes from selection on transmission rate: model

#### Methods

The initial selection analysis above clarifies how parasites and resources affect selection on resistance (quantified as transmission rate). However, in a dynamic system, evolution of this trait then feeds back onto densities of hosts, parasites, and resources. These eco-evo feedbacks are captured by evolutionary invasion analysis. In this analysis, fitness of an invading genotype with one resistance is evaluated in the environment created by a resident with another value of *β*_i_ (see eq. 3b for fitness). We determine invasion success of an invading genotype using a Next Generation Matrix proxy for fitness (Hurford et al. 2009; see Fig. A5 for pairwise invasibility plots). Then, we use Adaptive Dynamics (Eshel 1983) to find a Continuously Stable Strategy for resistance (*β*_CSS_). Near *β*_CSS_, genotypes with resistance closer to *β*_CSS_ can invade resident genotypes further from *β*_CSS_. Additionally, no genotype near *β*_CSS_ can invade a resident with *β*_i_ = *β*_CSS_ [see Brännström et al. (2013) for useful explanations and derivations]. Thus, *β*_CSS_ is a stable, eco-evolutionary attractor, one that incorporates feedbacks between resistance and density of resources and parasites (see Appendix section 1 for details).

Since host evolution now alters parasite density through feedbacks, we shifted focus to ecological drivers of epidemic size. Here, we chose carrying capacity of the resource (*K*; see Figs. A3, A4 for other drivers) because: higher *K* ecologically increases parasite density (Z); we manipulated *K* in mesocosms (below) by varying nutrient supply; and we find gradients of nutrient supply in nature that increase epidemic size (Fig. A6A). For a given value of *K*, we compute *β_CSS_*. Then, with that *β_CSS_*, we calculated infection prevalence and host density (at the stable equilibrium derived from eq. 1). Because biological disease outbreaks may not reach a stable equilibrium, we only use this equilibrium analysis to demonstrate biological logic and make qualitative predictions. Finally, we contextualize predicted prevalence and host density with purely ecological, single-genotype populations with fixed resistance (with values of *β*_i_ ranging from high to low, but without the possibility of evolution). More specifically, we calculate:

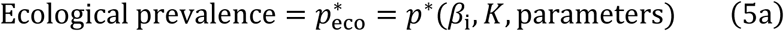

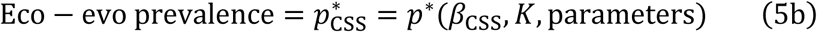

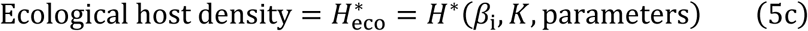

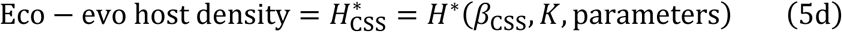

to determine how evolution of resistance affects prevalence and host density relative to meaningful, ecology-only controls. Although we varied resistance (with co-varying feeding rate; eq. 2) and carrying capacity, we held other parameters constant (Table 1). Then, we calculated equilibrium prevalence (*p\**_eco_ and *p\**_CSS_, eq. 5a, b) and host density (*H\**_eco_ and *H\**_CSS_; eq. 5c, d, respectively) for parasites which castrate (*θ* = 0) and those allowing full fecundity (*θ* = 1; see Appendix section 1 for outcomes with 0 < *θ* < 1). We found exactly one stable equilibrium in the parameter range for which we report results.

#### Results

As expected, evolution of increasing resistance (decreasing transmission rate) reduces prevalence and increases host density when parasites castrate. With castrating parasites (*θ* = 0), higher carrying capacity (*K*) increases density of parasite propagules (*Z*, not shown) when hosts cannot evolve (fixed *β*; Fig 2A); hence higher *K* increases prevalence (*p*\*; Fig 2B). In response, host populations evolve increased resistance [*β*_CSS_ drops with *K*; Fig. 2A], and prevalence (*p*\*_CSS_) flattens or even decreases carrying capacity (black curve; Fig. 2B). This pattern arises due to negative feedbacks between resistance and parasite density. In this feedback, increased *K* elevates parasite density, selecting for more resistance, thereby lowering parasite density (see also Fig. 3C, D). Notably, evolution of resistance enables hosts to attain increasing total (*H*\*_CSS_; Fig 2C) and especially susceptible [*S*\*_CSS_ = *m*/(*σβ*_CSS_); Fig. 2D] density as *K* increases (relative to fixed *β* controls). Hence, this castration case conforms to typical predictions of evolution of resistance during larger epidemics: resistance reduces prevalence and suppression of host density.

**Figure 3.**
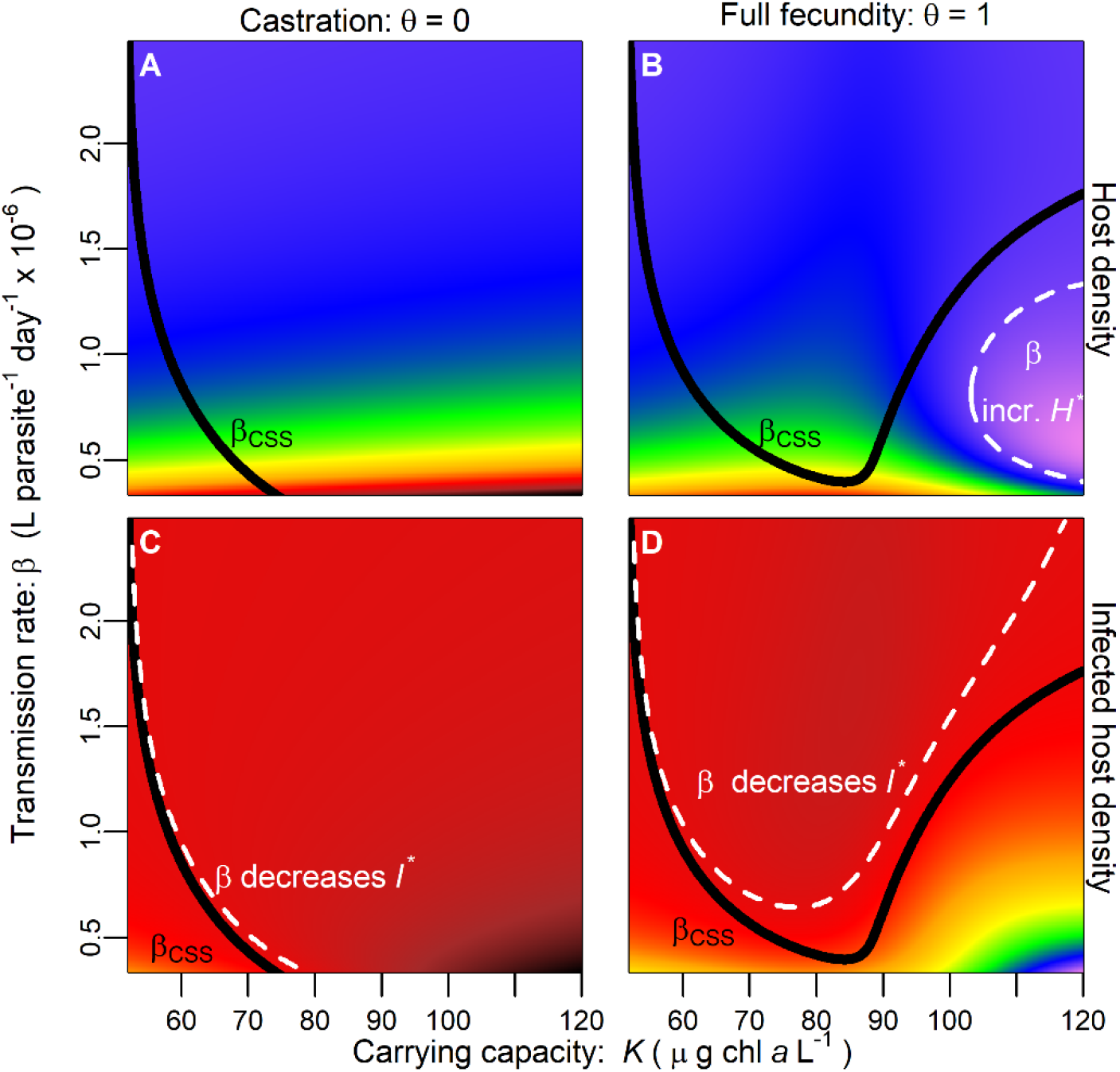
Evolution of lower resistance (higher β_CSS_) decreases host density but increases infected host density – an approach with ecological derivative maps. Color represents the effect of transmission rate, β_i_, on equilibrium host density [∂H*_eco_/∂β_i_; top row] or infected host density [∂I*_eco_/∂β_i_; top row], evaluated at a given combination of β_i_ and carrying capacity, K. Dark red denotes a strong negative effect of transmission rate on equilibrium density while light purple denotes a positive effect for both rows. (A) For castration (θ = 0), higher β_i_ decreases host density. The black curve demarks the β_CSS_ (see Fig. 2; it plunges below the axis range at c. K = 75). (B) For full fecundity (θ = 1), higher β_i_ usually decreases host density (∂H*_eco_ /∂β_i_ < 0). However, inside a small region (dashed white curve), higher β_i_ can increase it (∂H*_eco_ /∂β_i_ > 0). But the higher β_CSS_ does not enter this region (neither does lower β_CSS_: below y-axis range). (C, D) Infected host density, I, may increase with higher β_i_ (∂I*_eco_ /∂β_i_ > 0; below dashed white curve) or decrease if β_i_ is already too high (∂I*_eco_ /∂β_i_ < 0; above dashed white curve). The β_CSS_ values fall entirely inside the region of higher I with higher β (below dashed white divider), whether (C) with castration (θ = 0) or (D) full fecundity (θ = 1).

When parasites allow reproduction (e.g., shown with full fecundity [*θ* = 1]), the ‘resistance is futile’ response emerges with qualitatively different eco-evolutionary dynamics. At low carrying capacity (*K*), hosts evolve increasing resistance (lower transmission rate) with *K* (Fig. 2E). But when parasite density increases sufficiently (*p*\* _SS_ *≈* 0.4 and *K ≈* 85; Fig. 2F), the advantages of resistance decline (refer to Fig. 1). With even higher *K*, larger epidemics lead to evolution of decreasing resistance in a ‘resistance is futile’ effect. Now, positive feedbacks can arise between host evolution and parasite density; increased *K* elevates parasite density, selecting for less resistance (increasing portion of curve in Fig. 2E), which further elevates parasite density. These positive feedbacks are opposed by the stabilizing curvature of the tradeoff. But eventually the positive feedbacks become strong enough to create evolutionary bistability at higher *K*, with two CSS values of resistance possible (starting around *K* = 106; see Fig. A5 for pairwise invasibility plots). Evolution of decreasing resistance, then, increases prevalence. Carrying capacity increases prevalence ecologically (*p*\*_eco_; gray curves in Fig. 2F). At low *K*, evolution of increasing resistance slows (but does not prevent) an increase in prevalence (*p*\*_CSS_) with *K*. But at higher *K*, prevalence of the evolving host increases more quickly than for any ecological case. That prevalence pattern mirrors the response of the density of infected hosts (*I*\*) or parasite propagules (*Z*\*; both not shown but see Fig. 3D). Increased prevalence strongly depresses host density. Ecologically, host density increases with *K* (*H*\*_eco_; all gray curves). But evolution of less resistance lowers host density, potentially even causing it to decrease slightly with *K* (*H*\*_CSS_; slight dip in black curve Fig. 2G). Given the same host traits, a castrating parasite suppresses host density more (compare gray curves in Fig. 2C vs 2G) because it is more virulent. But ‘resistance is futile’ in the full fecundity case leads to lower eco-evolutionary host density than when parasites castrate (black curves in Fig. 2C vs. 2G), particularly for susceptible hosts (*S*\*, Fig. 2D vs. H).

As long as infected hosts retain some fecundity (0 < *θ* < 1), ‘resistance is futile’ arises at some point along gradients of carrying capacity (but at higher K with lower *θ*: Fig. A1). More broadly, ‘resistance is futile’ arises across other gradients (parameters) that would increase prevalence ecologically (e.g., higher parasite production, *σ*; Fig A3; lower loss rate of parasite propagules, *m*; Fig. A4). In these other instances, evolution of lower resistance enlarges epidemics and suppresses host density (Figs. A3, A4). The prevalence at which resistance starts becoming futile (*p*\*_CSS_ when *β*_CSS_ begins increasing with epidemic drivers) varies across a biologically feasible range (*p*\*_CSS_ = 0.42 in Fig. 2E-F, *p*\*_CSS_ = 0.24 in Fig. A3A-B, *p*\*_CSS_ = 0.049 Fig. A1A-B). Hence, ‘resistance is futile’ evolution – and the damage done to host density – may arise in a variety of biological scenarios with high drivers of epidemic size.

#### A complementary approach using ecological derivative maps

We more fully contextualize the ‘resistance is futile’ result using an ‘ecological derivative map’. In this map, we calculated the ecological effect of transmission rate (*β*, inverse of resistance), and thus also higher feeding rate [*f*_i_(*β*_i_)] on host density (*H*\*) in *β*-*K* (carrying capacity) space. For the castration case (*θ* = 0), higher *β*_i_ only decreases *H*\* (∂*H*_eco_*/∂*β*_i._< 0; Fig. 3A). But for full fecundity (*θ* = 1), higher *β*_i_ can increase *H*\* at high *K* and intermediate *β*_i_ (inside dashed white curve, ‘*β* incr. *H*\*’; ∂*H*_eco_*/∂*β*_i_ > 0; Fig. 3B). In this region, prevalence is high enough that increased *β*_i_ adds little additional mortality. Simultaneously, higher feeding rate can increase birth rate. The net balance can increase host density with *β*_i_ in this intermediate *β*_i_ range. Yet, with even higher *f*_i_ (and hence *β*_i_), further increases in *β*_i_ provide very little additional *f*_i_ (due to the curvature of the tradeoff) so higher *β*_i_ decreases host density. However, the CSS for transmission rate [*β*_css_] never enters the ‘*β* incr. *H*\*’ region (Fig. 3B). (The lower *β*_css_ attractor falls below the y-axis, where higher *β* only decreases *H*). Similarly, an ecological derivative map shows that higher *β*_i_ may increase or decrease density of infected hosts, *I*\*_eco_. The outcome depends on how *β*_i_ affects prevalence and total host density, as *I* = *p H*. However, higher *β*_i_ only increases *I*\*_eco_ in the evolutionarily relevant parameter space (outside of the ‘*β* decreases *I*\*’ region above the dashed white; Fig. 3C, D). Note that infected host and parasite propagule density are proportional, *Z*\* = *I*\**σ*(*δ*+*v*)/*m* from eq. 1d. Hence, for these parameter values, evolution of less resistance only reduces total density while increasing density of infected hosts.

### Eco-evolutionary outcomes from selection on resistance: experiment

#### Methods

We tested these predictions by creating experimental epidemics with a plankton system that meets many assumptions of the simple yet general model. The host, the zooplankton grazer, *Daphnia dentifera*, feeds on an alga, *Ankistrodesmus falcatus*. Carrying capacity of this resource depends on nutrient supply. Then, because hosts ingest infectious spores, they trade off feeding rate (and hence birth rate) with transmission rate of a fungal parasite, *Metschnikowia bicuspidata* (Auld et al. 2013; Hall et al. 2010; Hall et al. 2012). Hosts also vary in susceptibility to terminal infection once exposed (Stewart Merrill et al. 2019); this variation in susceptibility forms part of the tradeoff. Infected hosts retain significant fecundity, providing a test for the non-castration case (Ebert et al. 2000). In the 50 L mesocosms, thousands of individuals interacted dynamically with resources and parasites (as in the model) for approximately ten host generations.

We stocked mesocosms with two clonal genotypes of hosts with known (pre-measured) traits (Strauss et al. 2015). In monoclonal, ‘ecology-only’ reference mesocosms, we placed either a genotype with high resistance but slower feeding (hereafter: ‘high resistance genotype’) or one with lower resistance but higher feeding rate (‘low resistance genotype’; mimicking the gray curves in Fig. 2). In ‘evolving’ mesocosms, we stocked both genotypes at equal frequency, then allowed clonal frequency to change as clones competed during epidemics. These three host treatments were crossed with two nutrient levels (to mimic low and high *K*), yielding a 3 x 2 design with three replicates. Of these 18 populations, all but two outliers were included in the final analysis (see Appendix for more details). We sampled hosts twice a week to estimate density and infection prevalence. We used these samples of hosts twice during the epidemic for genotyping each evolving population (see Appendix section 2).

This experiment tested qualitative predictions of our eco-evolutionary model. Specifically, compared to populations in mesocosms that received lower supply of nutrients, high nutrient supply may experience ‘resistance is futile’ leading to larger epidemics, lower resistance, and lower host density. We tested the nutrients-prevalence prediction for the single-genotype populations with a beta regression of prevalence (because prevalence is bounded between 0 and 1). This model included nutrients, genotype, and their interaction (see Appendix section 2). We calculated mean transmission rate (*β*) for a treatment (Fig. 4B) or population (Figs 4C-F) as a weighted average of clonal estimates of that trait and genotype frequencies (estimated 16 and 24 days after spore addition; see Appendix section 2). We tested whether genotype frequency differed between nutrient supply treatments using a binomial regression of genotype identity on nutrients with mesocosm and time point as random effects. Then, we evaluated the relationship of transmission rate (x-axis) and prevalence (y-axis, beta regression) or host density (y-axis, total and infected, linear model) at each nutrient level (see Appendix section 2 for details).

**Figure 4.**
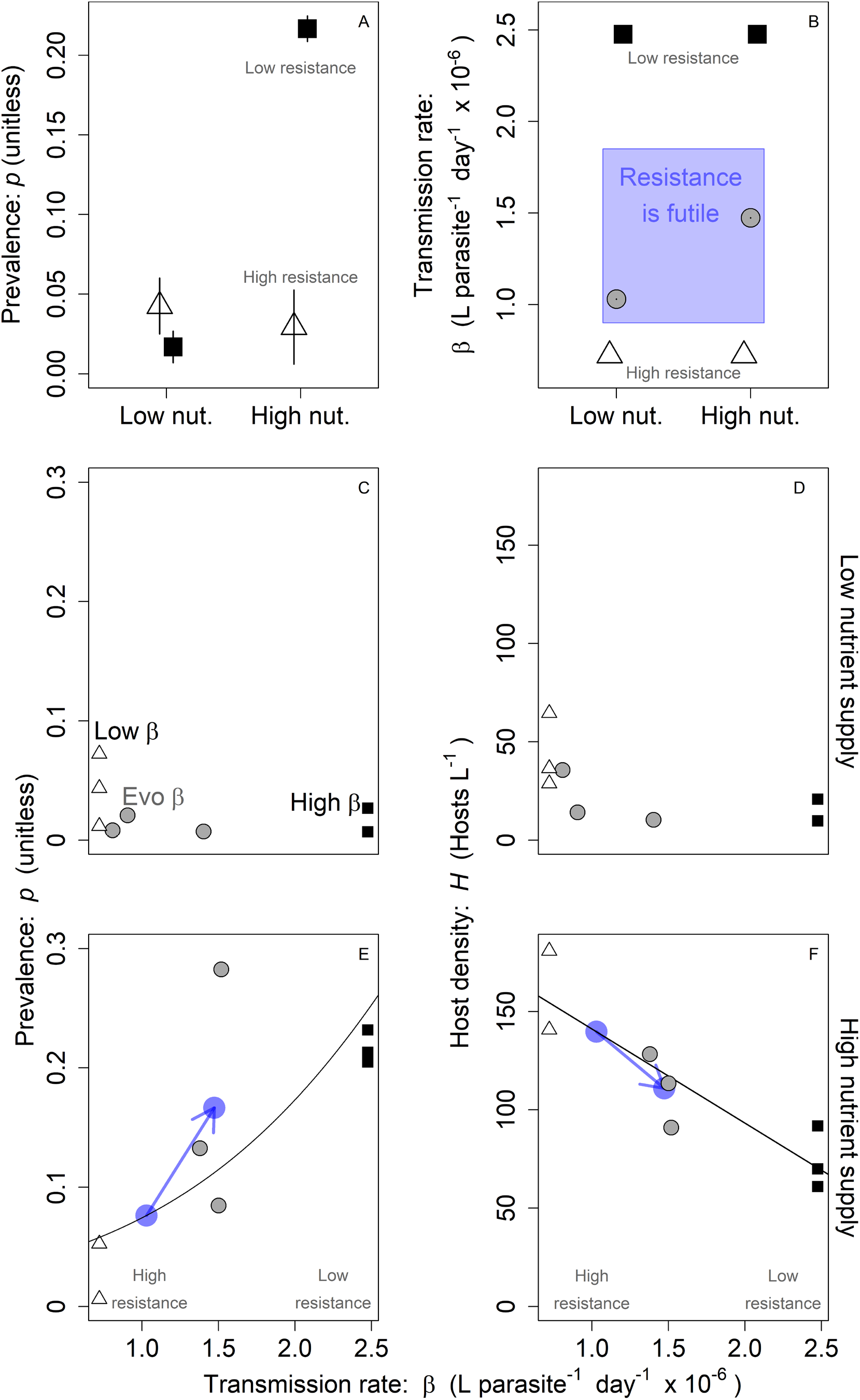
Implications of an epidemic driver (nutrient supply) for resistance, prevalence, and host density in populations that can evolve, relative to ecology-only controls. Symbols denote genotype treatments with c. three replicates each (two populations removed across entire experiment, e.g., due to extinction; see Appendix section 2). White triangles: monoculture of a high resistance genotype; black squares: a low resistance genotype; grey circles: both genotypes (potentially evolving via clonal turnover). (A) Ecology only: Prevalence (p) averaged over time and treatment for high or low resistance genotypes. Higher nutrients increase p for low resistance populations but not high resistance (significant overall effect of nutrients and interaction). (B) Resistance: High nutrient populations were less resistant, consistent with a ‘resistance is futile’ effect. Bars: ± 1 SE calculated over two time points and replicates within a treatment are barely visible. Low nutrient supply: (C) Prevalence stayed low as did (D) host density. High nutrient supply: Less resistance in (B) was associated with (E) higher p and (F) lower H. Blue arrows point from predicted values given resistance evolved at low nutrients to observed means at high nutrients.

#### Results

The experimental results qualitatively matched the eco-evolutionary model phenomenon of ‘resistance is futile’: less resistance evolved when epidemic drivers were high than when they were low. Resistance both decreased prevalence and decreased the positive effect of nutrients on prevalence (Fig. 4A; overall nutrient effect: P << 0.001; resistant genotype had lower prevalence: P = 0.0218; nutrient x genotype interaction P << 0.001). Then, in populations that could evolve, higher nutrient supply led to higher frequency of the low resistance genotype than at low supply (P = 0.0415 with 85 individuals IDed to genotype for this analysis; 17.5% less resistant genotype at low nutrients and 42.9% at high nutrients; see Fig. 4B). Thus, high nutrient populations did not evolve as much resistance (higher transmission rate, *β*) as low nutrient populations. At low nutrients, transmission rate did not correlate with prevalence (Fig. 4C; P = 0.389) or host density (*H*, Fig. 4D; P = 0.140). In contrast, at high nutrient supply, higher transmission rate correlated with higher prevalence (Fig. 4E; P = 0.0015) and lower host density (Fig. 4F; P = 0.0024). The eco-evolutionary model predicts that higher transmission rate should increase infected host density but this only trended non-significantly (not shown). Overall, the experiment largely matches the eco-evolutionary model’s qualitative predictions of ‘resistance is futile.’

The single-genotype reference populations contextualize the ecological effects of host evolution. If hosts at high nutrient supply evolved the resistance reached at low nutrients, the regression predicts they would have maintained lower prevalence and higher density (lower left blue point in Fig. 4E and upper left blue in Fig. 4F). Instead, high nutrient treatments had less resistance than low nutrient treatments (see Fig. 4B), presumably elevating prevalence (upper right blue in Fig. 4E) and suppressing host density (lower right blue: Fig. 4F). This eco-evolutionary result matches model predictions for ‘resistance is futile’ effects.

### Comparison of the distributions of epidemic size: models and the field

#### Methods

As we will show below, the model predicts that the ‘resistance is futile’ effect should make the distribution of epidemic size broader, somewhat bimodal, and right-skewed. Once we derived this result, we looked for it in the distribution of natural fungal epidemics. We sampled epidemics in Indiana lakes for 2009-2016 (166 unique lake-years). We measured prevalence of fungal infection multiple times over the course of the epidemic season (see Appendix: section 2 for details). We consider epidemic size in a given lake-year as maximum prevalence attained over the epidemic season (correlated with mean prevalence as shown in Fig. A6B; results for this section are similar if we use mean prevalence instead, see Table A1). We only consider lakes that had an epidemic (defined as reaching a maximum prevalence ≥ 0.1; varying this threshold did not change the key results of model competition: see Table A1). We compare this field distribution (Fig. 5A) to those produced from the eco-evolutionary model, given a distribution of the carrying capacity of the resource. (We chose *K* as the focal epidemic driver because it influenced prevalence in model and experiment, and likely in the field as well: see Fig. A6). Model distributions were generated under two cases of host evolution in the model with full fecundity while infected (*θ* = 1). In the ‘constrained’ case, we force hosts to evolve more resistance with *K*. To implement this constraint, we connect the decreasing portion of the upper eco-evolutionary attractor (Fig. 2E) to the lower one (with a tangent: Fig. 5B red curve). In the ‘resistance is futile’ case, hosts evolve normally, hence resistance increases then decreases with *K* (as in Fig. 2E, see Fig. 5B). Because of unmeasured factors in natural populations (e.g., other epidemic drivers), we did not expect a precise match between distributions of epidemic size in the model and field. Instead, we measured the deviations between them with descriptive statistics of breadth and shape: the interquartile range (IQR), bimodality (Ashman’s D ‘AshD’), and right-skew. A lower sum of relative deviations provided a better fit (found for each case using a genetic algorithm: see Appendix section 3).

**Figure 5.**
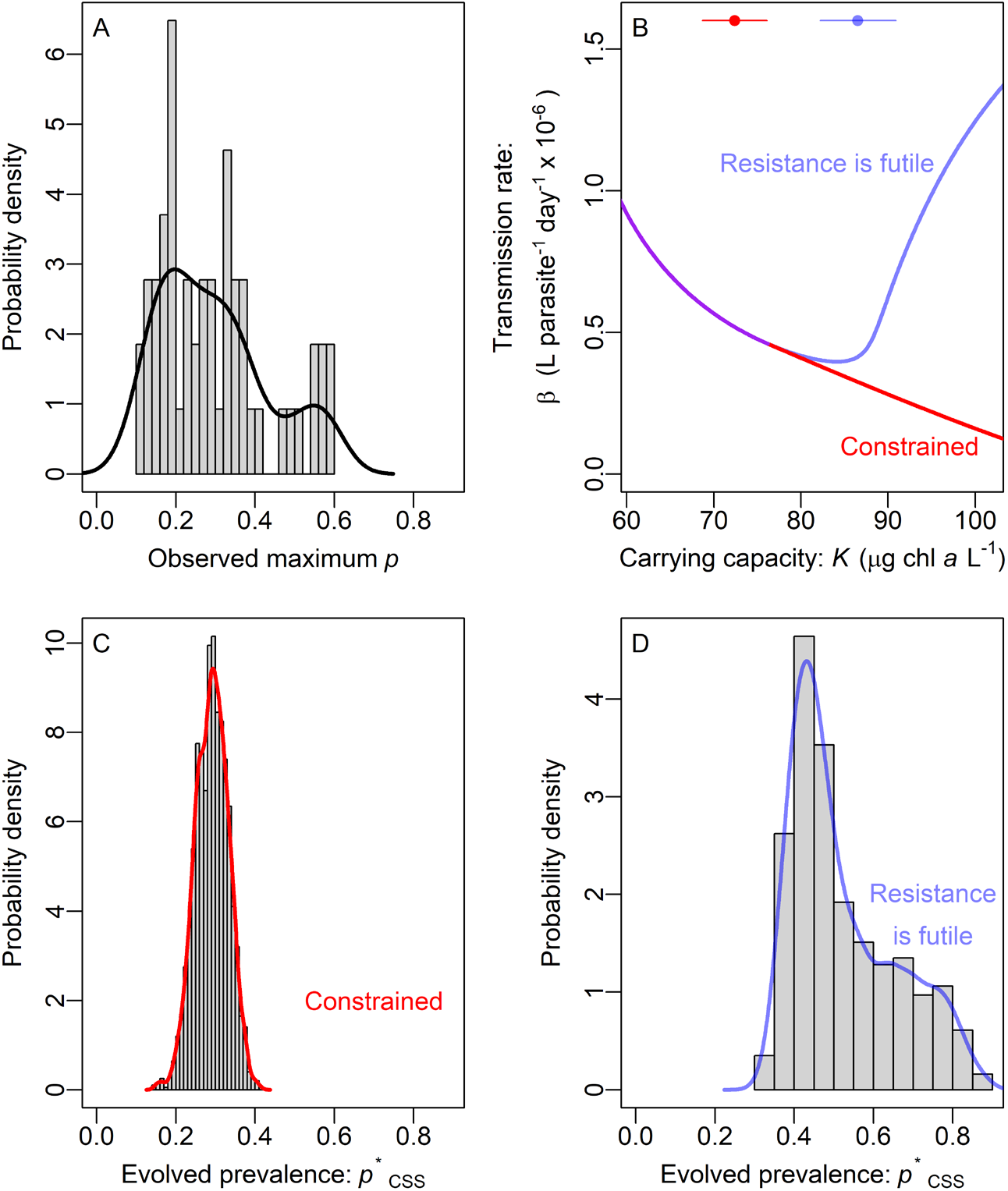
Comparison of model and natural epidemics. (A) Maximum prevalence in epidemics of the virulent fungus in Indiana (USA) lakes. Curves on histograms are smooth probability densities. (B) In the two model cases, hosts can evolve according to ‘resistance is futile’ (blue curve; compare Fig. 2E) or is constrained to only evolve increasing resistance with *K* (red curve). A purple curve is drawn where red and blue overlap. For each evolution case, we found the distribution of *K* yielding the best fit distribution of prevalence. Solid points and bars at the top of the plot indicate the mean ± sd *K* of the best fit distribution for each model case. (C) The ‘constrained’ eco-evolutionary model produced a narrow distribution of prevalence that matched the field data poorly. (D) The eco-evolutionary model with the ‘resistance is futile’ mechanism produced a broader, somewhat bimodal, right-skewed best fit distribution that better fit prevalence in Indiana lakes (compare to thin line, panel A).

#### Results

Natural populations displayed a broad, somewhat bimodal, right-skewed distribution of epidemic size. The observed distribution of epidemic size had IQR = 0.176, AshD = 3.33, Skew = 0.678 (Fig. 5A). Without ‘resistance is futile’ (Fig. 5B), host evolution creates a narrow (IQR = 0.0552, Fig. 5C), less bimodal (AshD = 1.43), and right-skewed (Skew = 0.00392) distribution that poorly matches the field data (summed relative deviance = 2.25). In contrast, the ‘resistance is futile’ mechanism produces a broader distribution (IQR = 0.208, Fig. 5F) with more bimodality and right-skew (AshD = 2.74, Skew = 0.696). These metrics better matched the field data (summed relative deviance = 0.361). The superiority of the ‘resistance is futile case’ holds for other analyses (i.e., lowering the epidemic size threshold and/or using mean prevalence: see Appendix section 3). In all instances, equilibria of the ‘resistance is futile’ case robustly produced larger IQR, AshD, and right-skew because of positive feedbacks between parasite density and host evolution. Hence, the broad, bimodal, right-skewed distribution observed in nature is more consistent with positive feedbacks that only arise in the ‘resistance is futile’ case of the eco-evolutionary model.

## Discussion

One might expect that hosts should evolve more resistance during larger epidemics of virulent parasites. However, as we argue here, resistance can be futile when epidemic drivers are high: more abundant, virulent parasites can select for less resistance to those parasites. That evolutionary response can increase infection prevalence and depress host density. We show this possibility in an eco-evolutionary model of host-parasite-resource dynamics. We confirm it qualitatively with a population-level experiment in a plankton system. Furthermore, the ‘resistance is futile’ response can create positive feedbacks between host evolution and parasite density. In the model these positive feedbacks produced broad, somewhat bimodal, right-skewed distributions of epidemic size consistent with those observed in natural epidemics. Hence, our theory, experimental results, and field survey all suggest that resistance can become futile when drivers of epidemics are high. That evolutionary response of hosts can increase prevalence and depress density of host populations.

Under what conditions do we expect to see the ‘resistance is futile’ effect? First, hosts must pay a cost for resistance, e.g., via a resistance-fecundity tradeoff. Second, low resistance genotypes must achieve higher fitness than high resistance genotypes when both become infected – a condition met when infected hosts retain (some) fecundity. Hence, ‘resistance is futile’ has not arisen when natural enemies eliminate future reproduction (e.g., castration) or mortality via predation (note these all consider resources as a driver of natural enemy abundance: Bohannan and Lenski 1997; Bohannan and Lenski 2000; Leibold 1996; Lopez-Pascua et al. 2014). Third, an epidemic driver must favor large enough epidemics to render resistance futile. To illustrate, our analyses (mathematical and empirical) focused on carrying capacity of the resource. Resource feedbacks (Buck and Ripple 2017) help to ensure that hosts survive such large epidemics (see Appendix section 1). How often do systems meet these conditions in nature? Various systems show the tradeoff (Auld et al. 2013; Boots and Begon 1993; Hall et al. 2010; Kraaijeveld and Godfray 1997); hosts often/usually retain some fecundity while infected (Kuris et al. 2008); and many systems have high prevalence of infection (Ezenwa 2004). A guppy-flatworm system likely possesses all three requirements (tradeoff: Hockley et al. 2014; Huizinga et al. 2009; reproduction: Pérez-Jvostov et al. 2015; very high prevalence in nature: Stephenson et al. 2015). When these conditions are met, larger epidemics can undermine selection for resistance.

Our model builds on an existing body of models demonstrating ‘resistance is futile’. Indeed, ‘resistance is futile’ arises during larger epidemics in other models, too (Best et al. 2017; Bonds et al. 2005; Donnelly et al. 2015; Miller et al. 2007) as does evolutionary bistability (Best et al. 2017; Miller et al. 2007). All of these models - despite differences in mechanisms – share a common thread. In each, high enough force of infection reduces the benefit of resistance (i.e., differential prevalence declines). Instead, reproduction by infected hosts rewards lower resistance (due to the fitness costs of resistance in a tradeoff). Eventually, the costs of resistance outweigh the diminishing benefits of resistance. However, our presentation extends beyond that previous work in three useful ways. First, we analyzed selection on fitness components. That separation produced clearer interpretation than approaches using fitness proxies (i.e., next generation matrices). Second, we showed the implications of ‘resistance is futile’ for prevalence and host density with comparison to ecology-only populations. That context stemmed from both ‘ecology-only’ controls and ‘ecological derivative maps’, two approaches that we advocate. Third, we showed why resources – an explicit ecological context – mattered for host evolution. We encourage others studying similar eco-(co)evolutionary problems to follow suit: delineate fitness components, characterize outcomes for prevalence and host density, and pinpoint the role of ecological context.

As importantly, we show direct experimental support for ‘resistance is futile.’ Specifically, the experiment saw hosts evolve less resistance during larger epidemics (created by higher nutrient supply) than during smaller epidemics. We found one other example (Parker 1991) of evolution of decreased resistance during a large epidemic but no examples of decreasing resistance with increasing epidemic size or epidemic drivers. In contrast, previous efforts showed evolution of higher resistance during larger epidemics (Duffy et al. 2012; Lopez-Pascua et al. 2014). Yet, our experimental demonstration matters because it showed how lower resistance can correlate with higher prevalence and lower host density. Hence, the ‘resistance is futile’ mechanism might explain some negative correlations between resistance and prevalence in nature (Ericson and Burdon 2009; Laine 2004; Thrall and Burdon 2000). We now have a second hypothesis that may contribute to such a pattern. The first was that negative correlations might emerge because less resistance causes higher prevalence. In the second, higher prevalence could cause lower (instead of higher) resistance when ‘resistance is futile’. Overall, our model-experiment combination challenges the general expectation that higher parasite density and or epidemic drivers should select for stronger resistance. They may not.

The implications of ‘resistance is futile’ for prevalence and host density raise broad, ecological concerns. When hosts evolve high resistance, infection prevalence remains small and epidemics depress host density little (Altizer et al. 2003; Christie and Searle 2018; Duffy and Sivars-Becker 2007). If hosts instead evolve less resistance during large epidemics, these epidemics may depress host density significantly. Furthermore, depressed density may threaten persistence of host populations (De Castro and Bolker 2005; Ebert et al. 2000). Additionally, ‘resistance is futile’ can produce high density of infected hosts (seen in the model, trended in the experiment). Higher infected density may increase risk of spillover to other host species (Daszak et al. 2000; Power and Mitchell 2004). Hence, evolution of lower resistance may exacerbate negative effects of large epidemics of virulent parasites for hosts of multiple species.

What are the similarities and differences between the ‘resistance is futile’ and evolution of tolerance? Both mechanisms maximize fitness while infected. Evolution of lower resistance (and higher fecundity) maximizes host fitness while infected. Similarly, tolerance may act by minimizing virulence on fecundity (Restif and Koella 2004) or mortality (Boots and Bowers 1999; Miller et al. 2006). Either the fecundity or mortality form of tolerance increases fitness of infected hosts (weakening virulence) without eliminating infection. Additionally, low resistance and tolerance both invoke positive feedbacks with parasites: higher parasite density can select for less resistance and higher tolerance (for tolerance, see Miller et al. 2005; Miller et al. 2006; Roy and Kirchner 2000). But the two mechanisms likely exert different consequences for host density. While lower resistance depresses host density, increased tolerance can elevate it (Miller et al. 2006). However, more complex possibilities arise: tolerance that lowers mortality may sometimes amplify directly-transmitted disease, thereby suppressing host density (Anderson 1979). The impact of tolerance on density will depend on epidemic size, costs of defense, and the precise trait value evolved. Just like our model of resistance, eco-evolutionary models of tolerance evolution (e.g., Best et al. 2017; Boots et al. 2009; Cressler et al. 2015) account for precisely these factors and could readily focus on prevalence and density in the future.

When will ‘resistance is futile’ arise, increasing prevalence and lowering host density, and when will it not? First, future tests of the eco-evolutionary model could employ a wider range and more levels of nutrient supply. Second, beyond nutrient supply, other factors should drive ‘resistance is futile’. Note, however, that direct manipulation of parasite supply would weaken feedbacks between host evolution and epidemic size. Third, different starting frequencies of genotypes could test for evolutionary bistability at high drivers of prevalence. Fourth, while laborious, trait measurements before and after epidemics (as in Duffy et al. 2012) could look for ‘resistance is futile’ during the largest epidemics (those driving and fueled by this mechanism). In addition, future theoretical work could alter key assumptions made here. First, if lower resistance increases mortality virulence (Hall et al. 2010), fitness advantages of low resistance strategies would shrink. Second, features of parasite evolution (e.g., important for Best et al. 2009; Slowinski et al. 2016) or dynamic parasite burden within hosts (Mideo et al. 2008) could be added. These extensions would highlight how and when increased size of epidemics leads to evolution of less resistance — and when it should not.

This study warns that evolution of lower resistance during larger epidemics may elevate prevalence and depress host density. This ‘resistance is futile’ outcome broadens the range of eco-evolutionary possibilities during epidemics. Even more importantly, ‘resistance is futile’ effects may increase prevalence and lower host density more commonly than anticipated. This possibility raises important implications for controlling the spread of disease. Yet, this eco-evolutionary mechanism requires several key components (fecundity-resistance tradeoffs, reproduction by infected hosts, and sufficiently large epidemics). At such an early stage, it remains unclear how many systems meet these assumptions. Guided by theory, we now know where to look: we can evaluate when maximization of fitness of individual hosts will amplify already-large epidemics and when/why it will not.

## Appendix

### Overview

We cover three major topics in this appendix. Section 1 presents further analysis of the model (eq. 1). We detail the computation of stable prevalence in fixed conditions. Further, we present modeling results on intermediate degrees of fecundity reduction (0 < θ < 1), the effects of and results for resources, and other parameter gradients. Lastly, we provide pairwise invasibility plots (PIPs) detailing the behavior of the evolutionary attractors. Section 2 provides more experimental and statistical detail for the mesocosm experiment. Section 3 describes observations of natural epidemics and comparison of statistical distributions to model epidemics.

### Section 1: Further analysis of the model

#### Determination of stable equilibria

The expression for model equilibria was found analytically in Matlab (Matlab R2017a) and exported to R (R Core Team 2019). For a given set of parameter values, we used linear stability analysis to determine which equilibrium was stable.

#### Computing stable prevalence for analysis of selection on transmission rate by parasites (Fig. 1)

Host fitness depends directly on prevalence of infection (eq. 3b). This prevalence determines the proportion of hosts suffering the virulent effects of parasites. In a constant environment, the density of resources (*R*) and parasites (*Z*) determine prevalence (*p*_i_) and thus fitness (*r*_i_). With this fitness, we can calculate derivatives of fitness with transmission rate (used to make Fig. 1). To calculate stable prevalence, we write the change in its prevalence through time. This prevalence depends on host traits (e.g., transmission rate, *β*_i_, and feeding rate, *f*_i_) and the environment (*R* and *Z*). If we define prevalence of genotype *i* as the proportion infected (i.e., *p*_i_ = *I*_i_ / [*I*_i_ + *S*_i_]), we can then write d*p*_i_/d*t* (eq. A1) and solve for stable prevalence, *p*_i stable_:

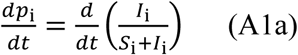

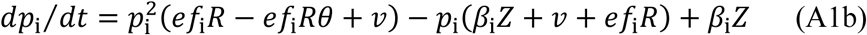

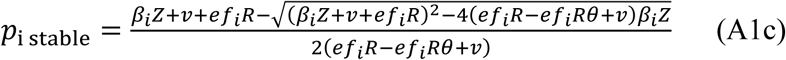

where *v* is added mortality virulence due to infection and *e* is conversion efficiency of births (as in Table 1). With this equation (eq. A1b, quadratic in *p*_i_ with fixed *R* and *Z*), we can solve for the stable prevalence (*p*_i stable_) that a genotype will suffer as a function of its traits, resources (*R*), and parasite propagules (*Z;* equ. A1c). This prevalence is stable (at d*p*_i_/d*t* = 0) because similar prevalence values move toward it 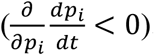. We numerically verified both stable prevalence and fitness found with this method using simulations (not shown). Further, we found (numerically) that it gave the same eco-evolutionary singular points as a more standard method using the Next Generation Matrix (comparison not shown). We use this stable prevalence method to calculate fitness directly for analysis of selection in a fixed environment. This method allows us to tease apart fitness into fecundity (*b*) and mortality (*d*). With that separation, we could then clearly see the effects of parasites on each fitness component (*b*, *d*). By calculating differential birth and death (∂*b*_i_/∂*β*_i_ and ∂*d*_i_/∂*β*_i_) components separately, we could better interpret response of fitness (*r*), hence selection to the resource-propagule environment.

#### Further explanation of Adaptive Dynamics to calculate full evolutionary outcomes (Figs. 2, 3)

Adaptive Dynamics finds a trait value that is an eco-evolutionary attractor called a “Continuously stable strategy.” Transmission rate at an eco-evolutionary attractor, *β*_CSS_, depends on competition between clonal genotypes with different transmission rates. This competition is modeled via evolutionary invasion analysis. In this approach, an invading genotype’s (‘a’) fitness, *r*_a inv b_, is calculated in the environment (resource and propagule density) created by a ‘resident’ genotype (‘b’). This quantity involves the invader’s transmission rate (*β*_a_) combined with the resource (*R*_b_) and propagule (Z_b_) environment set by the resident. For the resident to be a CSS, genotypes with similar transmission rate must have negative fitness when invading (mathematically, this requirement means that ∂*r_a inv b_*/∂*β_a_* = 0 and ∂^2^*r_a inv b_*/∂*β_a_^2^* < 0). Those conditions guarantee that the *β*_CSS_ cannot be invaded. Then, host populations must evolve toward that transmission rate *β*_CSS_. Such ‘convergence stability’ requires that genotypes closer to *β*_CSS_ can invade genotypes further from *β*_CSS_; the fitness gradient (∂*r_a inv b_*/∂*β_a_*) must be positive when *β*_b_ < *β*_CSS_ and negative when *β*_b_ > *β*_CSS_ [mathematically, d/d*β*_b_ (∂*r*_a inv b_/∂*β*_a_) < 0]. We use the Next Generation Matrix method (Hurford et al. 2009) to calculate the matrix’s dominant eigenvalue (giving the basic reproductive number, *R*_0_, i.e., expected lifetime reproduction of a host) as a proxy for fitness. While this proxy is not fitness, it is positive when invader fitness is positive and negative when invader fitness is negative. Thus, NGM gives the same solution for *β*_CSS_. All derivatives are evaluated at *β*_a_ = *β*_b_ = *β*_CSS_ (Eshel 1983). Also see Brännström et al. (2013) for useful explanations and derivations. Thus, we find the eco-evolutionarily stable transmission rate (*β*_CSS_) given feedbacks between transmission rate and density of resources and parasites.

#### Outcomes from fecundity reduction (extension of Fig. 2)

When parasites reduce but do not eliminate fecundity (0 < *θ* < 1), eco-evolutionary outcomes qualitatively resemble the outcomes when the parasite does not alter fecundity (*θ* = 1) along a carrying capacity gradient. When fecundity reduction is less severe (Fig. A1A, solid curve, *θ* = 0.9), hosts can evolve increased transmission rate (higher *β*, decreased resistance) at relatively low *K*. When fecundity reduction is more severe (Fig. A1A, dotted curve, *θ* = 0.3), *K* must reach higher levels for hosts to evolve higher *β*. That level of *K* may correspond to fairly low prevalence (*p*_CSS_; i.e., less than 10% for *θ* = 0.3; Fig. A1B). But once hosts begin to evolve higher transmission rate, prevalence can quickly increase (e.g., *p*\*_CSS_ sharply increases to almost 1 for *θ* = 0.3) — despite that parasites exert strong costs to host fitness. High prevalence of virulent parasites then supresses host density (i.e., the higher *β_CSS_* branch leads to higher *p*\*_CSS_ producing the lower branch of *H*\*_CSS_; Fig. A1C). Hence, hosts can evolve high *β* if infected hosts retain some fecundity (*θ* > 0). Without castration, genotypes with high *β*_i_ retain a fecundity advantage through the feeding rate-transmission rate tradeoff (eq. 2). And given high enough carrying capacity, differential prevalence shrinks so that the fecundity advantage can lead to a ‘resistance is futile’ effect, large epidemics, and large host declines (Fig. A1).

**Figure A1.**
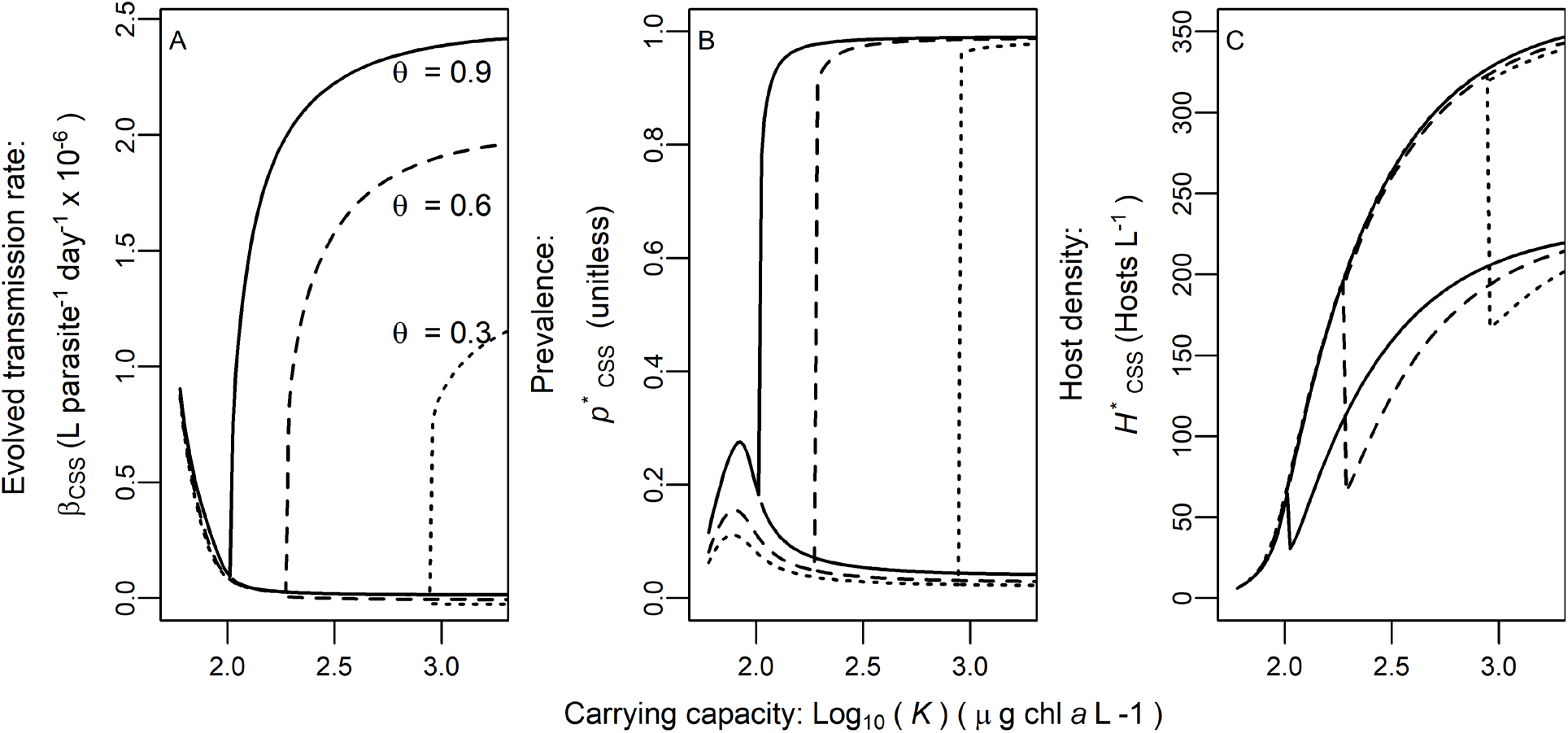
Outcomes from fecundity reduction (compare to Fig. 2). When parasites reduce but do not eliminate fecundity (0 < *θ* <1), a ‘resistance is futile’ effect arises along a carrying capacity gradient (*K* shown on log_10_ scale) as in the no effect of infection on fecundity case (*θ* = 1; Fig. 2E-H). (A) At low *K*, hosts evolve decreasing transmission rate (low *β_CSS_*). Then, hosts can evolve decreasing (low *β_CSS_* curve) or increasing transmission rate (high *β_CSS_* curve) at high *K*. Evolutionary repellers between alternative *β_CSS_* curves are not shown to minimize visual clutter. When parasites reduce fecundity more strongly (e.g. dotted curve *θ* = 0.3), increasing *β* evolves at higher *K*. (B) When hosts evolve decreasing transmission rate, prevalence (*p*\*_CSS_) remains low (at the lower *β*_CSS_ attractor). If hosts evolve to high *β*_CSS_, *p*\*_CSS_ increases strongly. Increasing *β*_CSS_ can begin at *p*\*_CSS_ = 0.22, 0.049, or 0.024 (for *θ* = 0.9, 0.6, or 0.3 respectively) (C) Host density (*H*\*_CSS_ = *S*\*_CSS_ + *I*\*_CSS_) increases with *K* but is greatly reduced when hosts evolve to the higher *β*_CSS_ attractor. Host density declines even more sharply for susceptible host density (not shown here but similar to Fig. 2H). Some curves shifted slightly vertically for clarity.

#### The role and effects of resources

Resource feedbacks help hosts survive sufficiently high prevalence of infection to enable ‘resistance is futile’. This response arises while infected hosts can reproduce (*θ* > 0). An extreme case clearly illustrates this point (100% prevalence: derived from eq. 3b for *p*_i_ = 1):

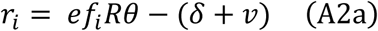

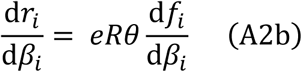

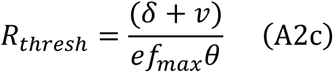

Fitness of these completely infected hosts (*r*_i_, eq. A2a) is fecundity while infected (*ef_i_Rθ*) minus mortality enhanced by mortality virulence (*δ*+*v*). Since *p*_i_ = 1, higher transmission rate (*β*_i_) leads to higher fitness (*r*_i_) if *θ* > 0 (since the *f*-*β* tradeoff ensures d*f*_i_/d*β*_i_ > 0; eq. A2b). So, reproduction by infected hosts guarantees that a ‘resistance is futile’ effect will eventually arise as prevalence increases (at the very least, for the asymptotic case of *p* = 1). This effect can also emerge at less extreme values of prevalence (e.g., Figs. 1D, 2F, A1B) depending on the tradeoff strength (d*f*_i_/d*β*_i_) and other parameters. The key to the ‘resistance is futile’ effect, then, is that differential prevalence (d*p*_i_/d*β*_i_) declines with higher prevalence (dotted curve declining in Fig. 1A). This decline must occur eventually because d*p*_i_/d*β*_i_ = 0 at *p*_i_ = 1 – that forces differential prevalence to peak when 0 < *p_i_* < 1. However, note that this extreme case requires that the host population can survive 100% prevalence. Such survival requires that resources exceed a minimum threshold (*R*_thresh_, eq. A2c) set by the maximum feeding rate (*f*_max_). While a ‘resistance is futile’ effect generally does not require *p*_i_ = 1, sufficient resources are still required to support hosts.

This resource threshold in the analysis of selection hints that, in the fully eco-evolutionary model, resource feedbacks promote survival at high prevalence and thus help enable a ‘resistance is futile’ effect. Parasites kill hosts and/or reduce fecundity; thus, during epidemics, the minimal resource requirement of hosts increases, elevating resource density (Buck and Ripple 2017). These increased resources, in turn, more likely satisfy the survival condition (*R*_thresh_ for *p*_i_ = 1; more complex for *p*_i_ < 1; see below). Thus, feedbacks which elevate resource density promote a ‘resistance is futile’ effect. Host evolution, in turn, feeds back onto resource density (see Fig. A2). Hence, full eco-evolutionary outcomes depend on feedbacks between evolution and ecology (e.g., resource and parasite density) of hosts.

For a ‘resistance is futile’ effect to arise at prevalence less than one, a more complex resource threshold must be exceeded. We derive (from eq. 3b) the general derivative of fitness with respect to transmission rate (d*r*_i_/d*β*_i_):

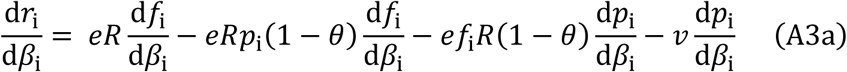

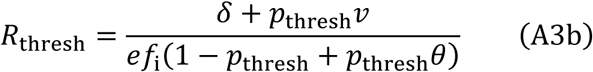

If infected hosts reproduce (*θ* > 0), this selection gradient (eq. A3a) will eventually become positive once *p*_i_ crosses some threshold of prevalence, *p*_thresh_ (see Figs. 1D and A1A; it collapses to eq. A2b at *p*_i_ = 1). But hosts may not survive with prevalence this high. Persistence at *p*_thresh_ requires a threshold density of resources (eq. A3b, which collapses to eq. A2c at *p*_thresh_ = 1; for *p*_thresh_ < 1, *f*_i_ is some feeding rate and may be smaller than *f*_max_ since *f*_max_ is only guaranteed to be selected for at *p*_i_ = 0 or 1). So, during epidemics, parasites indirectly elevate resources. If this resource response is large enough, hosts can persist during epidemics that trigger a ‘resistance is futile’ effect.

Host evolution, in turn, impacts resource density. Equilibrium resource density depends on host mortality divided by fecundity per resource: *R*\* = (*δ* + *vp*) / [*ef*(1 – *p* + *θp*)]. A population having both higher transmission rate (*β*) and feeding rate (*f*) via the tradeoff will suffer higher mortality from disease (due to higher prevalence, *p*). Generally, this higher mortality will indirectly drive higher resource density (*R*\*). This pattern holds for castration (*θ* = 0; Fig. A2A). It mostly holds without castration but there are exceptions (dashed gray rises higher than solid gray in Fig. A2B). Here, *R*\* is maximized at intermediate *β* when *K* is high. This pattern arises because increasing *β* only slightly increases prevalence and population-level mortality when prevalence is already very high. More importantly, higher *β* is tied to increasing feeding rate (*f*). So, when hosts are already highly infected, higher *β* has little effect on *p* (numerator of *R*\*) but increases *f* (denominator of *R*\*).

Higher resource density also strengthens selection for higher feeding rate (eq. A3a). We determine the importance of changing resource density (impacting differential fecundity) compared to changing differential prevalence (impacting differential mortality). We calculate differential fecundity and differential mortality at a representative *β*_1_ (1 x 10^-6^); we find qualitatively very similar results at all other relevant values of *β*. The constant values of parasite and resource density used correspond to CSS outcomes at a given *K*, i.e., *Z*\*_CSS_ and *R*\*_CSS_. When differential fecundity is greater than differential mortality, the selection gradient (differential fitness) is positive, indicating selection for *β*_CSS_ > *β*_1_. When differential mortality is greater, selection favors a lower transmission rate. When differential fecundity and mortality are equal (intersections in Fig. A2C), *β*_CSS_ = *β*_1_ (*β*_1_ also meets other CSS criteria not shown). Over the range of *K*, differential mortality responds to *K* much more strongly than differential fecundity. Thus, these parameters indicate that changing differential prevalence is more important for driving ‘resistance is futile’ than changing resource density is.

**Figure A2.**
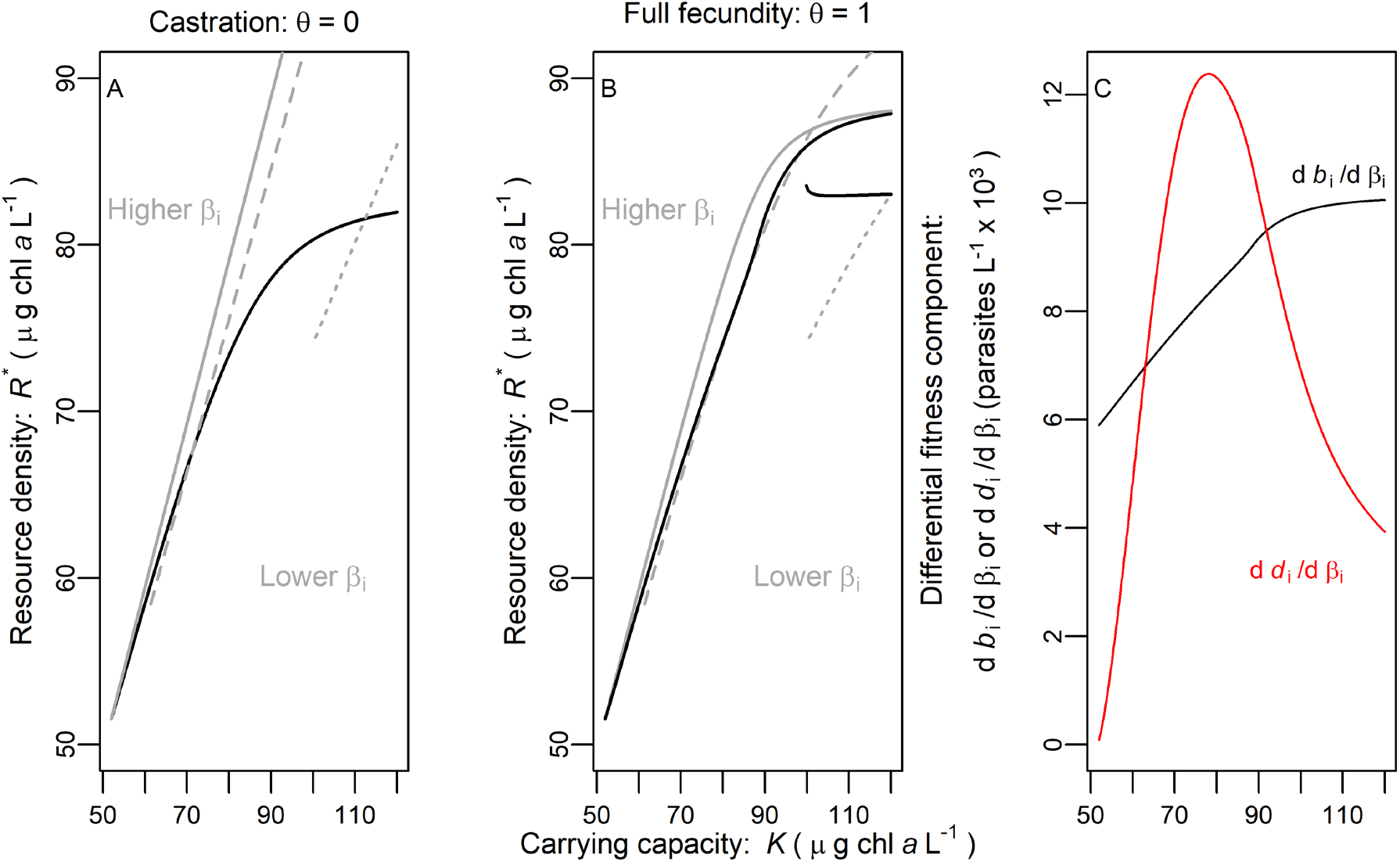
Feedbacks of host evolution and resource density. (A-B) Black curves are resource density at the eco-evolutionary attractor [*R*\*_CSS_]. Gray curves are single-genotype reference populations [*R*\*_eco_] with the same values of transmission rate (*β*_i_) shown previously (Fig. 2). Gray curves only drawn in the range where the endemic equilibrium is feasible. (A) For castration (*θ* = 0), resource density increases with carrying capacity (*K*: gray curves increase for higher [solid], intermediate [dashed], and lower [dotted] values of transmission rate). Higher transmission rate (lower resistance) also leads to higher resources. With higher carrying capacity, hosts evolve increasing resistance [decreasing *β*_CSS_] so that resources increase more slowly than for the single-genotype cases. (B) For no effect of infection on fecundity (*θ* = 1) resource outcomes are more complex. Higher values of transmission rate do not always increase resources in the single genotype references (e.g., gray lines cross). There are also two eco-evolutionary attractors (as in Fig. 2E-H). The lower transmission rate attractor corresponds to lower resource density. (C) Differential fecundity (black curve) and differential mortality (red curve) are shown in an environment fixed at *Z*\*_CSS_ and *R*\*_CSS_ (along higher *β*_CSS_ attractor). Differential mortality responds to *K* more strongly than differential fecundity does. Shown for *β*_1_ = 7.96 x 10^-7^ and qualitatively similar for other *β*.

#### Eco-evolutionary outcomes along other gradients

Evolution of less resistance arises at high prevalence of infection, not just for high carrying capacity (*K*). We arrived at this conclusion by varying parameters that increase or decrease parasites ecologically. Besides *K* (Figs. 2, A1), other parameters increase ecological prevalence of disease (and density of parasite propagules): conversion efficiency (*e*), maximal growth rate of resources (*w*), or parasites released per infected host (*σ*). As those parameters increase, hosts eventually evolve less resistance in a ‘resistance is futile’ effect (moving to the right; Fig. A3) and bistability arises. Other traits (parameters) decrease parasite propagule density: background mortality rate of hosts (*d*), loss rate of parasites (*m*), or mortality virulence (*v*). As these traits decrease (moving to the left in Fig. A4), hosts evolve more resistance until a point. At that point, hosts may evolve less resistance, promoting prevalence and harming host density. If instead hosts evolve to an evolutionary attractor with higher resistance, then prevalence remains low and host density reaches a higher level. Therefore, gradients of a variety of traits that promote parasite density can produce a ‘resistance is futile’ effect.

**Figure A3.**
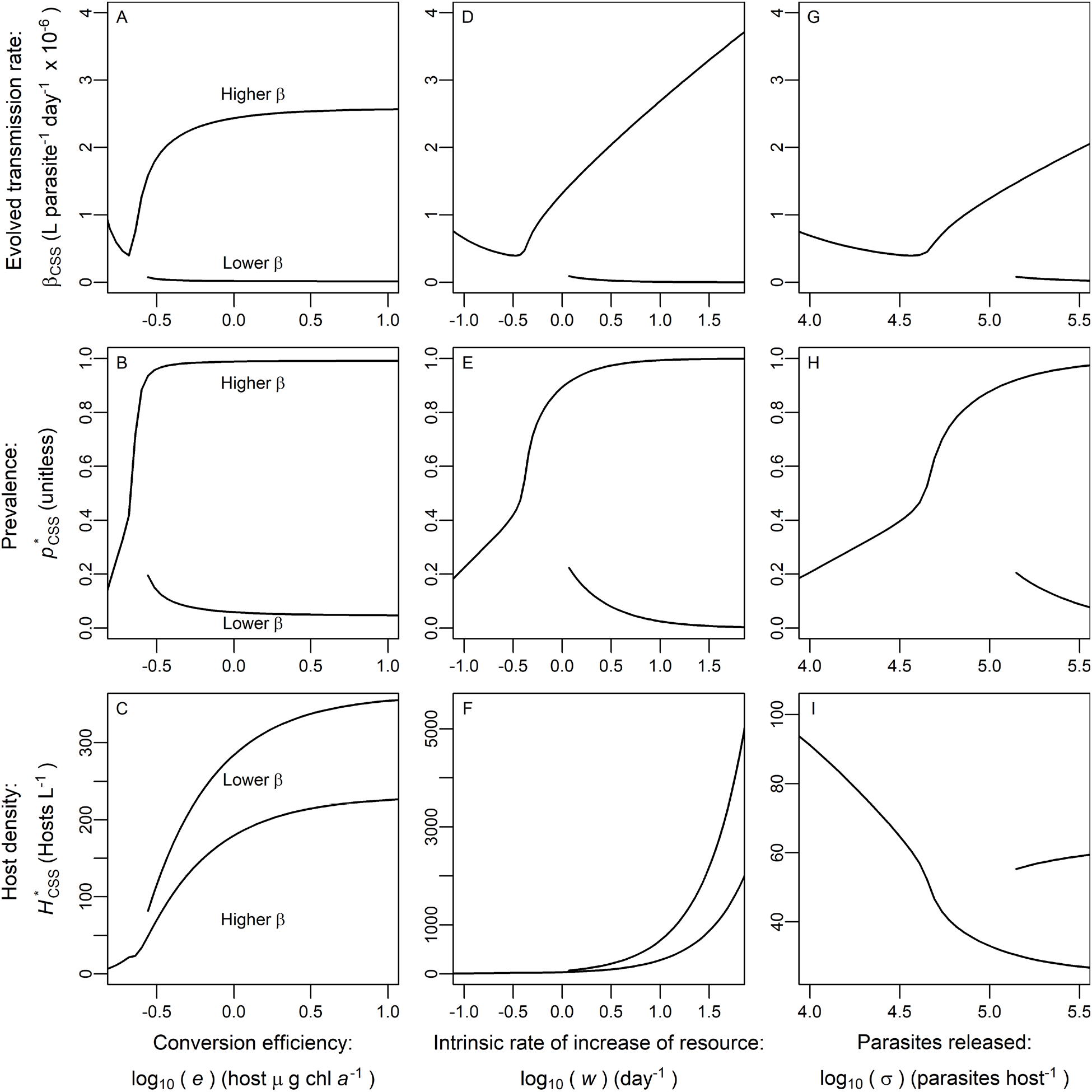
Eco-evolutionary effects of parameters that (ecologically) increase disease (extension of Figs. 2, A1). X-axes on log_10_ scale. Increases in several parameters besides carrying efficiency of hosts (*e*; panels A-C), resource intrinsic rate of increase (*w*; D-F), and parasites released per infected host (*σ*; G-I). (A) Increasing conversion efficiency ecologically increases density of parasite propagules (for ecology-only populations). More parasites first increase differential prevalence, selecting for lower transmission rate (*β*_CSS_ curve decreases). Then increasing parasites decreases differential prevalence, selecting for higher transmission due to fecundity benefits (the ‘resistance is futile’ effect; *β*_CSS_ curve begins to increase; *θ* = 1 shown). Lastly, alternative eco-evolutionary attractors (two *β*_CSS_ curves) can arise. Evolution of higher transmission rate leads to (B) high prevalence and (C) lower density (especially density of susceptible hosts; not shown). Lower transmission rate leads to lower prevalence and higher host density. This reasoning applies for *w* (D-F) or *σ* (G-I). Default parameter values are as in Table 1.

**Figure A4.**
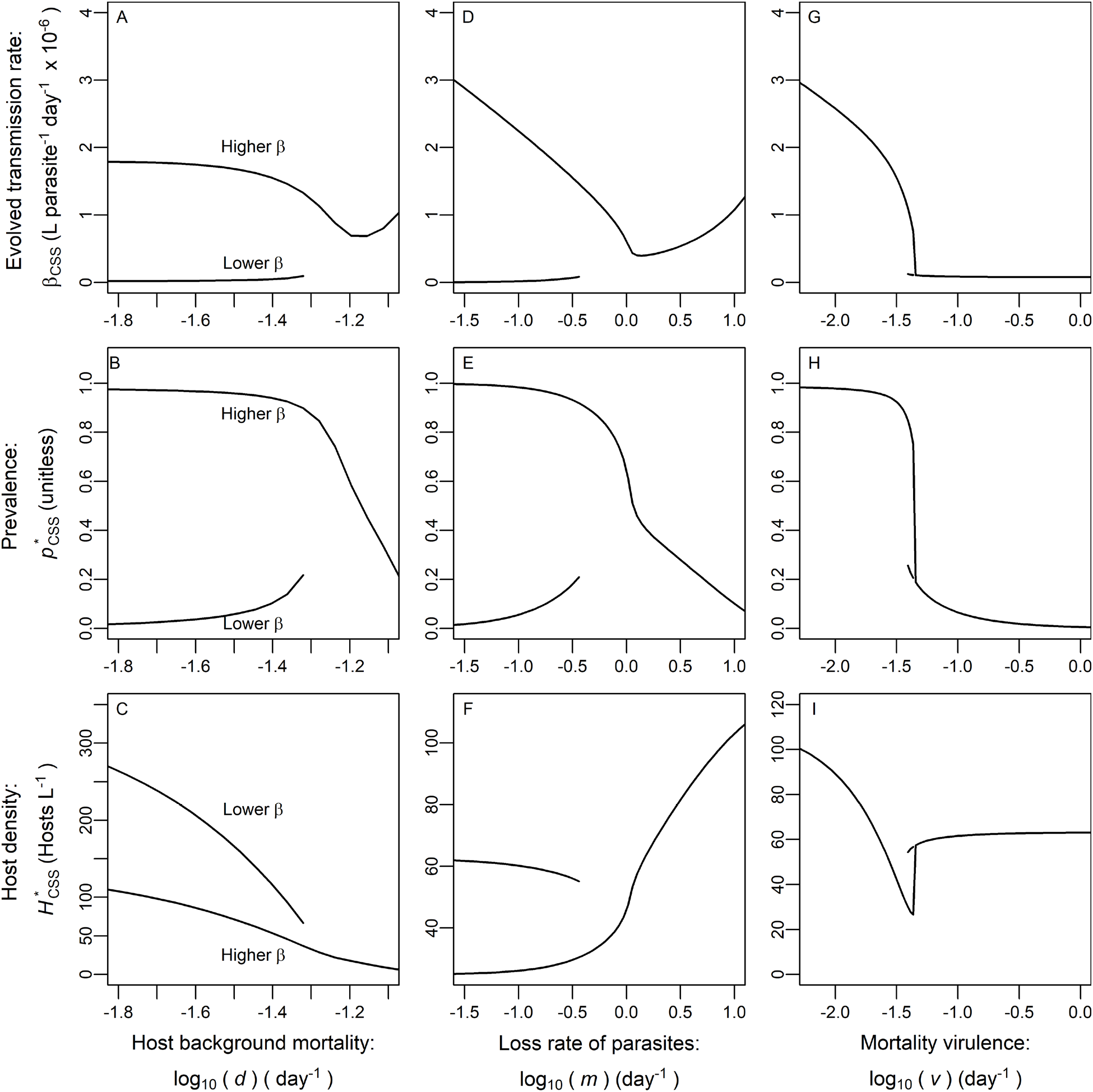
Eco-evolutionary effects of parameters that (ecologically) decrease disease (extension of Figs. 2, A1, A3). X-axes on log_10_ scale. Increases in several parameters decrease epidemics: host background mortality (*d*; panels A-C), loss rate of parasites (*m*; D-F), and mortality virulence (*v*; G-I). Reading each panel from right to left (i.e., increasing disease prevalence), the reasoning follows that in Fig. A3 left to right (also increasing disease prevalence). (A) Decreasing background mortality of hosts (*d*) ecologically increases parasite density (for ecology-only populations). More parasites select for decreasing transmission rate (*β*_CSS_ curve decreases moving right to left) until fecundity benefits of higher transmission reverses selection (the ‘resistance is futile’ effect; *β*_CSS_ curve increase to the left; *θ* = 1 shown). Then, alternative eco-evolutionary attractors (two *β*_CSS_ curves) can arise. High transmission rate leads to (B) high prevalence and (C) lower host density (especially for susceptible hosts, not shown). Low transmission rate leads to low prevalence and higher host density. Similar reasoning applies for *m* (D-F). The results are somewhat different for *v* (G-I). Default parameter values as in Table 1.

Eco-evolutionary patterns deviate with mortality virulence of parasites (*v*). Ecologically, decreasing *v* increases prevalence, parasite density, and host density. The evolutionary attractor with lower transmission rate is only stable over a small range of *v* values (Fig. A4G). When infection is almost avirulent (*v* is low), there is little advantage to avoiding infection. Further, when infection becomes harmful at high *v*, hosts evolve low transmission rate despite lower prevalence. Harm to host density is maximized at intermediate *v* where prevalence is somewhat high and infection is somewhat deadly (Fig. A4H, I). This means eco-evolutionary host density can actually decline with decreasing *v*, unlike ecological host density (note these outcomes would differ for a directly transmitted parasite instead of the obligate-killing, environmental parasite we model). Hence, along a gradient of *v*, eco-evolutionary feedbacks can have opposite effects on host density than purely ecological ones.

Evolution of very low transmission rate can invert the ecological effect of several other gradients of traits (parameters). Evolution of very low transmission rate in the castration case (*θ* = 0) or lower CSS (when *θ* > 0) decreases prevalence along gradients of carrying capacity (*K*), conversion efficiency (*e*), intrinsic rate of increase of resources (*w*), and parasite production per host (*σ*). Without evolution (i.e., for the reference genotype cases with purely ecological feedbacks), increases in these parameters (*K*, *e*, *w*, *σ*) would increase prevalence (Fig. 2B, F; Figs. A1B, E, H). Similarly, when background mortality of hosts (*d*) or loss rate of parasites (*m*) decrease, prevalence would increase ecologically. But, evolution of hosts to a very low *β*_CSS_ causes prevalence to decrease (Fig. A4B, E). Additionally, host density follows a similar pattern. Evolution of decreased transmission rate can increase host density when it would otherwise decrease ecologically. Increasing production of parasite propagules (*σ*; Fig. A3I) or mortality virulence (*v*; Fig. A4I) decrease host density ecologically. But evolution to the lower *β*_CSS_ increases host density along these gradients. Thus, when hosts evolve low transmission rate, prevalence and density of hosts respond oppositely to gradients with full eco-evolutionary feedbacks (*p*\*_CSS_, *H*\*_CSS_) than with strictly ecological feedbacks (*p*\*_eco_, *H*\*_eco_).

#### Pairwise invasibility plots

**Figure A5.**
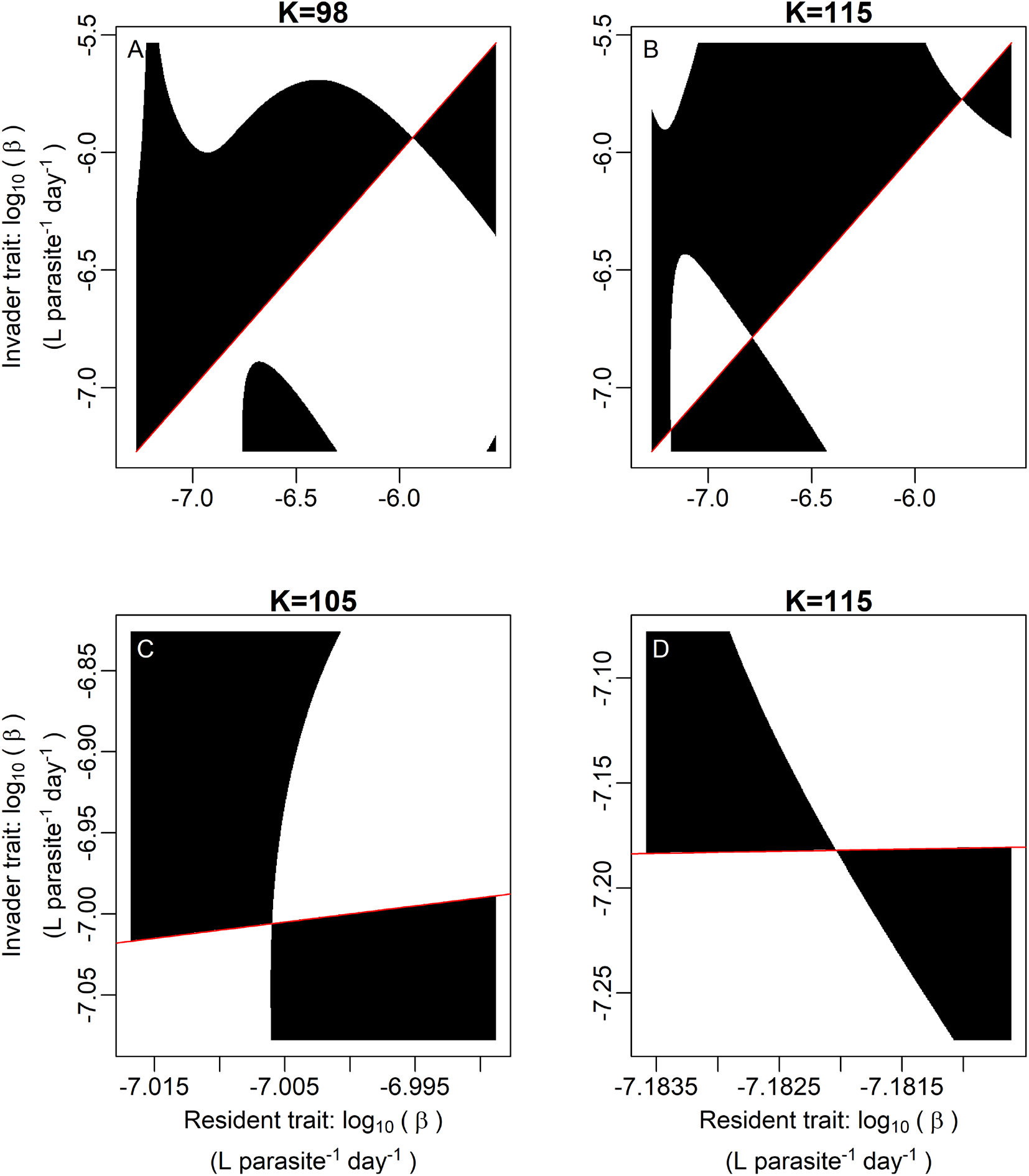
Pairwise invasibility plot shows how evolutionary singular strategies (β_i_ such that selection gradient = 0) change stability properties. Black regions denote positive invader fitness and white negative fitness. The 1 to 1 line is shown in red for clarity, especially since the clearest axis ranges are not always 1 to 1. Other parameters follow defaults (Table 1). (A) At low K, there is only one evolutionarily singular strategy and it is a CSS (as in Fig. 2E). (B) At higher K, two new singular strategies appear. One is a repeller between the lower and higher CSS. Zooming in on the lowest singular strategy, which changes stability behavior: (C) When it first appears, the lowest singular strategy is convergence stable but invadable by nearby strategies, i.e., an evolutionary branching point. (D) Once K is somewhat higher, the lowest singular strategy is no longer invadable by nearby strategies and is a CSS.

### Section 2: Further methods and results for the mesocosm experiment

#### Methods for mesocosm experiment

Each mesocosm was housed in a 75-liter acid washed polyethylene tank held in a climate-controlled room at approximately 21°C. We filled tanks to 50 L with 80% tap water (treated with Kordon Amquel Plus and Novaqua plus) and 20% filtered (Pall A/E: 1 µm) lake water. We added water to compensate for evaporative losses as needed. Additions of nitrogen (as NaNO_3_) and phosphorus (as K_2_HPO_4_) were added to low supply (5 μg L^-1^ P) and high supply (50 μg L^-1^ P) levels; nitrogen was added at 20:1 N:P ratio by mass. Nutrients were replenished twice weekly throughout the experiment to account for an estimated (exponential) 5% per day loss rate. All tanks were inoculated with 50 mg (by dry weight) of the green alga *Ankistrodesmus falcatus* 7 days before hosts were introduced (algae day −6, hosts day 1) to reach sufficient density to support hosts.

We added hosts to each tank on day 1. Monocultures were stocked at 10 hosts L^-1^; the evolution treatment started with a 50:50 ratio of the two genotypes, 5 hosts L^-1^ of the high resistance (‘Bristol 10’) genotype and 5 hosts L^-1^ of the low resistance (‘A4-3’) genotype. Once introduced, hosts then grew 27 days before addition of 3,600 fungal spores L^-1^ (day 28). Prior to introduction into the mesocosms, the two isoclonal host lines were obtained from Midwestern (MI) USA lakes and cultured in the laboratory while spores (from Baker Lake, Barry Co, MI, USA) were cultured *in vivo* by passage through live hosts. Tanks received a 16 L: 8 D light cycle after host addition. Twice a week on days 14-86, we sampled 1 L of tank water, then sieved animals through 80 µm mesh to destructively sample hosts. We visually counted and diagnosed hosts for infection using dissecting microscopes (40-50X).

Population averages were taken over a 28-day time window (days 48 – 76) to best estimate equilibrial quantities predicted by our mathematical models. On day 48, most populations (inoculated day 28) began to display sufficient numbers of visible infections. We ended on day 76 because mesocosms accumulated detritus and dynamics became less consistent across replicates. Then, over this 28-day window, we calculated averages as area under the curve divided by time. One low-nutrient, low resistance genotype alone replicate was excluded from the analysis because that population went extinct early in the experiment. One high-nutrient, high resistance genotype alone replicate was excluded as a severe outlier with extremely low host density, possibly due to chemical contamination. This population was the only population with a Cook’s distance > 4X the mean for host density (corresponding to 95^th^ percentile) and more than twice the Cook’s distance of any other population; such a deviation is uncharacteristically low for this genotype, even at lower nutrient supplies (personal observation). Each estimate of an individual population’s prevalence and host density included at least 84 total hosts sampled over the experiment with the average being 525.3. These samples represented a fraction of the population with the smallest population having an average size of 489 individuals.

#### Measuring host trait evolution

Given that key traits of individual hosts were measured previously (Strauss et al. 2015), trait evolution could be determined from clonal frequency. The experimental methods to estimate traits are explained in detail there (Strauss et al. 2015). In brief here: feeding rate for each host genotype was measured via declining algal concentration in tubes containing single animals. Transmission rate was determined from the probability of infection given a specific spore dose (either 100 or 450 spores mL^-1^) and duration of exposure (22 hours). Our two genotypes traded off transmission rate with feeding rate, allowing for evolution along a tradeoff (*β*_1_ = 7.22 x 10^-7^ L parasite^-1^ day^-1^, *β*_2_ = 2.48 x 10^-6^, *f*_1_ = 0.0124 L host^-1^ day^-1^, *f*_2_ = 0.0227). We determined average transmission rate in evolving host populations using clonal frequencies.

We sampled clonal genotype frequencies twice for each evolving population. We preserved up to 10 hosts (if present in 1 L sample) from each evolving tank in 70% ethanol with 5% 0.5 mM EDTA, then stored them at 2°C for later genotyping. We genotyped samples collected from 16 days (day 44) and 34 days (day 62) after fungal inoculation. DNA preservation, extraction, amplification, and identification of genotypes via microsatellite loci followed previously published methods (Strauss et al. 2017) at the Roy J. Carver Biotechnology Center at the University of Illinois Urbana Champaign. Weighted mean transmission rate was calculated from previous trait measurements (above) and genotype frequencies pooled from both time points (up to 20 animals) for the purposes of Fig. 4B. As mentioned in Methods, the effect of nutrient supply on host evolution was determined from a binomial regression of genotype identity on nutrients.

#### Statistical methods for the mesocosm experiment

We tested key model predictions regarding nutrients, prevalence, host evolution, and hosts density using mesocosm data. Because prevalence is bounded between 0 and 1, we regressed prevalence upon nutrient and single-genotype treatment (Fig. 4A) with a beta regression (Ferrari and Cribari-Neto 2004; Mangiafico 2016). Higher nutrients significantly increased prevalence overall in single-genotype populations (P-values reported in main text). In a significant interaction, nutrients increased prevalence more for the high transmission rate genotype. Diagnostic plots (following Ferrari and Cribari-Neto 2004) supported the validity of the beta regression’s assumptions. Thus, nutrients ecologically increased prevalence, providing a gradient for testing host evolution. We tested whether hosts evolved higher transmission rate (in two-genotype populations) at higher nutrients (Fig. 4B) with a mixed effects binomial regression using the lme4 package in R. Time (day 44 or 72) and replicate were included as random effects. The low resistance genotype was more frequent at high nutrients than at low nutrients. Thus, hosts evolved less resistance when ecological drivers of prevalence were higher.

We then tested the effects of transmission rate on prevalence, host density and infected host density. For prevalence, we used beta regression; diagnostic plots (as above) met model assumptions. For host and infected host densities, we used linear models for which diagnostic plots supported assumptions of the linear model (normality, homoscedasticity, linearity). Thus, evolution of higher transmission rate increased prevalence and decreased host density.

### Section 3: Distribution of natural epidemics is consistent with positive feedbacks from a ‘resistance is futile’ effect

#### Overview

It was logistically infeasible to estimate transmission rate among many clones in the lake-years of the field survey, particularly since ‘resistance is futile’ would not be expected to occur in all lake-years. Hence, we took an indirect approach: we compared the observed distribution of the size of natural epidemics to distributions generated by the eco-evolutionary model of a parasite that allows full fecundity. We sampled the size of natural epidemics in Indiana lakes from 2009 to 2016 (166 unique lake-years). When a ‘resistance is futile’ effect is impossible in the model, evolution of higher resistance with more parasites lowers parasite prevalence (and thus lower density of parasite propagules). Thus, there arises a net negative feedback loop between resistance evolution and density of parasite propagules. These negative feedbacks alter the distribution of epidemic size, reducing variance (measured by interquartile range) and also affecting bimodality and skew. When a ‘resistance is futile’ effect is possible, hosts can evolve decreasing resistance with increasing parasites. Because lower resistance increases parasite density, positive feedbacks arise. These positive feedbacks alter the distribution of epidemic size, increasing interquartile range, bimodality, and right-skew. We independently optimize the fits of model distributions of epidemic size with and without ‘resistance is futile’ evolution to the distribution observed in natural populations and compare summed relative deviance. We create variation in model epidemic size arising from variation in carrying capacity. To evaluate the relevance of this driver, we analyze the effect of total phosphorus on natural epidemic size as well (assuming this measure provided an estimate of carrying capacity).

#### Distribution of natural epidemics: Methods

We sampled natural fungal epidemics *Daphnia* hosts in lakes in Greene and Sullivan counties, Indiana from 2009-2016. Individual lakes were sampled in the following years: Airline (2009-11, 2013-16), Beaver Dam (2009-16), Benefiel (2009-11, 2014-16), Canvasback (2009-15), Chapel (2015-16), Clear (2015-16), Corky (2014-16), Dogwood (2009-11, 2014-2016), Downing (2009-16), Frank (2009, 2014-16), Front (2015-16), Gambill (2009-16), Goodman (2009-2016), Goose (2009-11, 2014-2016), Hackberry (2014-2016), Hale (2009-10, 2014-16), Horseshoe (2015-16), Island (2009-15), Long (2009-16), Lonnie (2014-15), Mayfield (2009-11, 2014), Midland (2009-16), Narrow (2014-16), Pump (2009-16), Scott (2009-16), Shake 1 (2014-16), Shake 2 (2015-16), Shop (2014), Spencer (2015-16), Star (2015-16), Sycamore (2014-16), T Lake (2009-10, 2014-16), Todd (2009-11, 2014-16), Trout (2014-15), Walnut (2015-16), Wampler (2015-16), Willow (2014). Thus, 166 individual lake-years were sampled.

One lake-year gave one data point for this analysis. Lakes were sampled at least every other week September through November (epidemic season for the fungal parasite). In some years, sampling extended outside this window (e.g., July 11 - Dec. 5 in 2011) and/or was conducted weekly (weekly in 2011). Each sampling, one vertical net tow was taken from each of three different sites within the deep basin of the lake using a 13 cm diameter, 153-μm mesh Wisconsin bucket net. These samples were pooled and diagnosed (200 individuals or greater) under a dissecting microscope to determine the proportion of animals suffering visible infection. Many lake-years did not experience appreciable epidemics. For example, in 47 lake-years no infected animals were detected and 27 other lake-years had some infected individuals but never reached 1% infection. Thus, we considered lake-years that *did* experience sizable epidemics (i.e., maximum prevalence had to reach at least 10%; robustness to this threshold tested below). The mean prevalence over the course of an epidemic was calculated as the average area under the curve starting at the first date when infection was observed in a year to the last. Maximum and mean prevalence were closely related (see below). Hence, both provide appropriate comparison to model equilibria. Seasonal factors (e.g., temperature Shocket et al. 2018) strongly affect the spread of disease and timing of epidemics. Thus, prevalence averaged over time approximates a changing eco-evolutionary attractor. Alternatively, maximum prevalence approximates the eco-evolutionary attractor when it is positioned to most spread disease. Both measures are characterized.

#### Distribution of natural epidemics: Results

Natural populations displayed a highly variable, somewhat bimodal, right-skewed distribution of epidemic size. We characterize distribution shape with interquartile range, Ashman’s D, and skewness. There is no single consensus on the best summary statistic for bimodality; hence, we use Ashman’s D as a reasonable choice. This measure of bimodality (*AshD*) was determined from means and standard deviations (SD) of two underlying Gaussian distributions (Ashman et al. 1994):

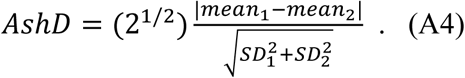

Values of Ashman’s D (eq. A4) which exceed 2 are necessary to clearly separate the underlying distributions if they are Gaussian (Shi et al. 2017). For our measure of bimodality, we fit a Gaussian mixture model (2 underlying distributions) to prevalence (maximum or mean) using the mclust package in Rstudio (R Core Team 2019). We found skewness of distributions using the desctools package in Rstudio (R Core Team 2019). The observed distribution of maximum prevalence (Fig. 5A) had IQR = 0.176, AshD = 3.33, Skew = 0.678. Results were similar for mean prevalence (see Table A1). These distributions may indicate underlying positive feedbacks between host resistance evolution and parasite density in natural populations.

#### Distribution of model epidemics: Methods

We compare how well model distributions can fit this natural distribution. For each model, we generate prevalence distributions by allowing one parameter to vary lognormally with a certain mean and standard deviation (SD). We varied one parameter for simplicity; to maintain consistency with the mesocosm experiment, we chose carrying capacity of the resource (*K*). We randomly sample the distribution of *K* (2000 values) and then find the value(s) of transmission rate to which hosts evolve for each value of *K* [*β*_CSS_, Fig. 5B]. With *β*_CSS_ and *K*, we find prevalence [*p*\*_CSS_ (*β*_CSS_, *K*); all other parameters as in Table 1]. Thus, we map a lognormal distribution of *K* to find a distribution of prevalence. Then, we find IQR, Ashman’s D, and skew for the resulting model distribution. We summed relative deviance between the model (M) distribution and observed (O) distribution for all three measures: summed relative deviance = |IQR_O_-IQR_M_ |/ IQR_O_ + |AshD_O_-AshD_M_| / AshD_O_ + |Skew_O_-Skew_M_| / Skew_O_ (where |…| indicates absolute value). Lower summed relative deviance indicates better fit.

We optimize this fit for each model independently with a genetic algorithm. Genetic algorithms apply flexibly to many optimization problems because they make few assumptions (Whitley 1994). In each ‘generation’ of the genetic algorithm, we find eco-evolutionary prevalence distributions from 50 distributions of carrying capacity, *K* (mean and SD). The best fit in a generation is the prevalence distribution with the lowest summed relative deviance from the observed distribution. Whichever has lower deviance between this best fit and the best fit of all previous generations is passed on to the next generation. The mean and SD of this winning distribution are passed to the next generation and slight mutations to it produce 49 new *K* distributions that also populate the next generation of the algorithm. Mutation of mean or SD occur by multiplying them by 10^x_i_ (where x_i_ is the i^th^ independent observation of a normally distributed random variable with mean = 0 and SD = 0.001). Generation 0 is initialized with the same distribution of *K* for both models (untransformed mean = 81.6, SD = 4.07). We ran the algorithm for generations 1-100 independently for the model cases with and without ‘resistance is futile.’ Summed relative deviance over generations of the algorithm (not shown) indicated the number of possible distributions per generation (50) and generations (100) were sufficient for asymptotic approach to an optimal fit of the model to the field data. Then we determined which model cases’ best fit most closely matched the observed distribution of prevalence.

#### Distribution of model epidemics: Results

Positive feedbacks in the ‘resistance is futile’ case make it better fit the observed distribution of prevalence. When hosts evolve more resistance during larger epidemics, these feedbacks can only be negative. For heuristic benefit, we visualized the effect of evolution on prevalence (and thus parasite density) by comparing eco-evolutionary prevalence (*p*\*_CSS_) to ecology-only prevalence (*p*\*_eco_). We calculated ecology-only prevalence with an arbitrary transmission rate (*β*) that gives *p*\*_eco_ that overlaps with *p*\*_CSS_. This choice of *β*_i_ for *p*\*_eco_ is purely cosmetic; it does not alter whether negative feedbacks (negative slope of *p*\*_eco_ vs *p*\*_CSS_ -*p*\*_eco_) or positive feedbacks (positive slope) can arise. Both *p*\*_eco_ and *p*\*_CSS_ used the best fit *K* distribution (lognormal, untransformed mean = 86.7, SD = 4.38 with ‘resistance is futile’ and mean = 72.4, SD = 3.70 without) selected by the genetic algorithm to fit the observed distribution of epidemic size. Without ‘resistance is futile’, increasing *K* increases *p*\*_eco_; meanwhile, hosts evolve lower and lower *β*_CSS_ leading to a more negative effect of host evolution on prevalence (Fig. 5C). This curve only has negative slope, representing negative feedbacks of parasite density (tied to prevalence) on evolution of transmission rate of hosts. The model case with only negative feedbacks did not fit the field data as well as the model case with negative and positive feedbacks.

We varied threshold prevalence and the use of maximum or average prevalence to test the robustness of this analysis. We crossed the use of maximum (default) or average prevalence with requiring maximum prevalence to reach 0.10 (default) or 0.01 for a lake-year to be included in our analysis. The results are shown in table A1. For all four combinations, the model case with ‘resistance is futile’ fits better than the one without because of higher interquartile range, bimodality, and right-skew.

**Table A1.** Robustness of model-field comparison to prevalence metric and threshold. Epidemics must reach a maximum prevalence of at least 0.1 or 0.01 (depending on version of analysis) to be included in the analysis (*p* threshold). The metric of field prevalence used may be either maximum prevalence over an epidemic season or mean prevalence over time (*p* metric). Statistics are reported for the field data and each model case. “RiF” and “Constrained” are the model cases with and without ‘resistance is futile.’ The model with the lower summed relative deviance (“Deviance”) is the better fit.

| $p$ threshold | $p$ metric | Statistic | Data | RiF | Constrained |
| --- | --- | --- | --- | --- | --- |
| 0.1 | Max | IQR | 0.176 | 0.208 | 0.0552 |
| 0.1 | Max | AshD | 3.33 | 2.74 | 1.43 |
| 0.1 | Max | Skew | 0.678 | 0.696 | 0.00392 |
| 0.1 | Max | Deviance | 0 | 0.386 | 2.25 |
| 0.1 | Mean | IQR | 0.0732 | 0.116 | 0.0577 |
| 0.1 | Mean | AshD | 2.88 | 2.35 | 1.48 |
| 0.1 | Mean | Skew | 0.719 | 1.27 | 0.0768 |
| 0.1 | Mean | Deviance | 0 | 1.37 | 1.61 |
| 0.01 | Max | IQR | 0.248 | 0.192 | 0.0555 |
| 0.01 | Max | AshD | 2.13 | 2.27 | 1.61 |
| 0.01 | Max | Skew | 0.818 | 0.761 | -0.0135 |
| 0.01 | Max | Deviance | 0 | 0.361 | 2.04 |
| 0.01 | Mean | IQR | 0.0841 | 0.0951 | 0.0578 |
| 0.01 | Mean | AshD | 1.95 | 1.91 | 1.76 |
| 0.01 | Mean | Skew | 0.867 | 1.44 | -0.0460 |
| 0.01 | Mean | Deviance | 0 | 0.811 | 1.47 |

#### Relationships between total phosphorus and prevalence

Total phosphorus content, a putative index of carrying capacity of the resource (*K*), correlates with increased prevalence in natural populations. Epilimnetic total phosphorus was analyzed by colorimetric spectrophotometry using the ascorbic acid method following persulfate digestion (Prepas and Rigler 1982). Average infection prevalence was found as detailed below while maximum prevalence was self-explanatory. Because prevalence is bounded between 0 and 1, we fit a beta regression to prevalence with lake and year treated as random factors. Diagnostic plots (i.e. QQ and residual vs. predicted plots) from the DHARMa package in Rstudio (R Core Team 2019) indicate key assumptions of this statistical test were upheld. Average phosphorus correlates with increased prevalence (Fig. A6A). Average prevalence and maximum prevalence could be used to characterize the distribution of epidemic size and are closely correlated (beta regression; average prevalence as independent variable; Fig. A6B). Varying the required size of epidemics (max prevalence > 0.1 or 0.01) did not qualitatively affect these outcomes.

**Figure A6.**
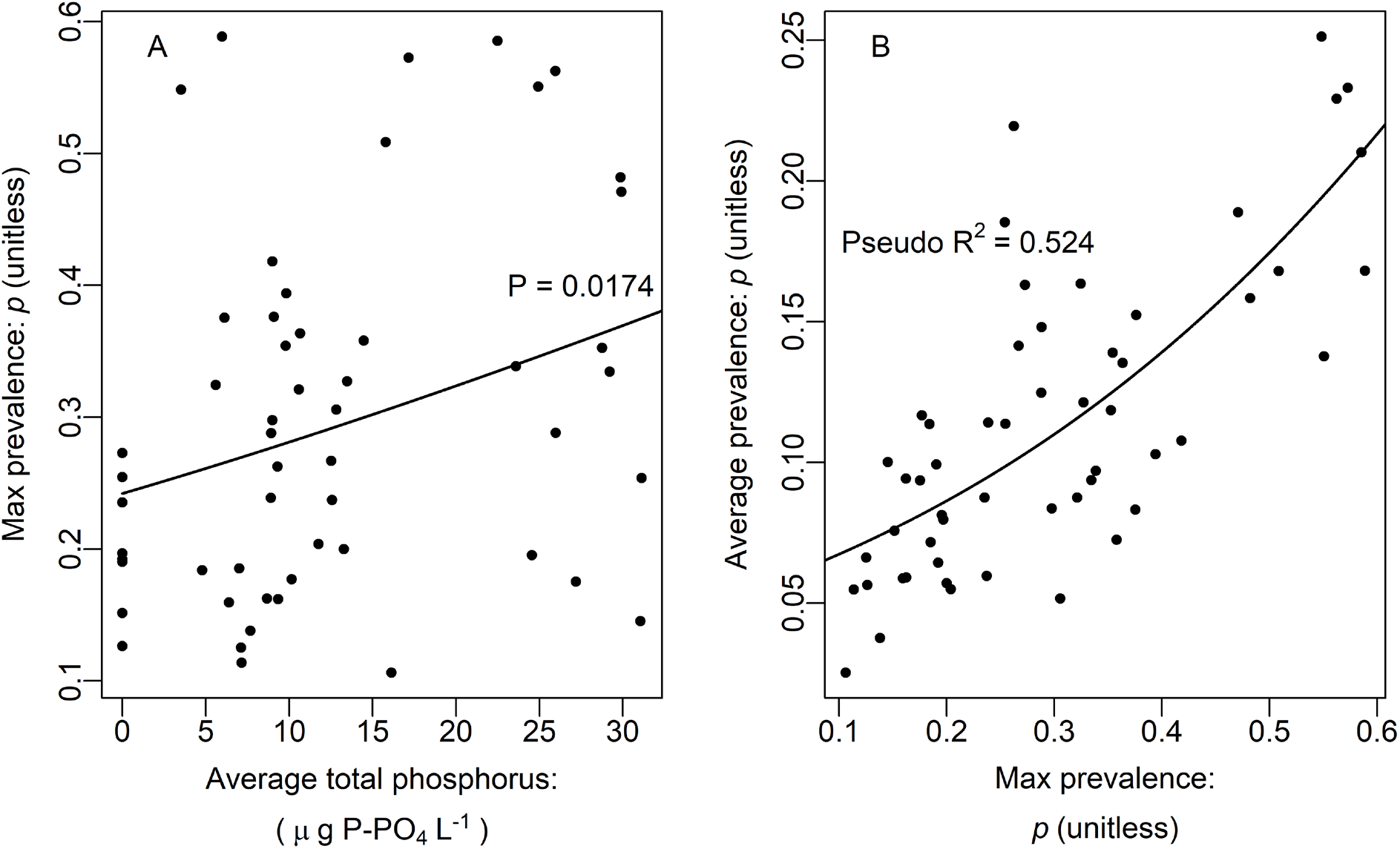
Prevalence in natural epidemics of fungal parasites in the zooplankton hosts. Indiana lakes were sampled repeatedly during fungal epidemics over multiple years. Each point represents one lake in one year. Average total phosphorus and average prevalence (proportion of hosts infected) were calculated as area under curve divided by time of epidemic duration. Duration of epidemics lasted from the first time during epidemic season that infected individuals are observed until the last time. A given lake-year is considered as a significant epidemic if maximum prevalence was at least 0.10 (10% of hosts infected). These trends remained, qualitatively, for a smaller prevalence threshold (0.01). Prevalence was fit by beta regressions (see text) with lake and year as random effects. (a) Prevalence of infection increases across a gradient of total phosphorus. (b) Average prevalence correlated with maximum prevalence among lake years.

